# Numerical Simulation of Transdermal Delivery of Drug Nanocarriers Using Solid Microneedles

**DOI:** 10.1101/2023.12.27.573440

**Authors:** Ben Newell, Wenbo Zhan

**Affiliations:** School of Engineering, University of Aberdeen, Aberdeen, AB24 3UE, United Kingdom

**Author notes:** Correspondence: Dr Wenbo Zhan, School of Engineering, Old Aberdeen Campus, University of Aberdeen, Aberdeen AB24 3UE, UK.

**Keywords:** Mathematical model, Medicated adhesive patch, Nanocarrier, Solid Microneedle, Transdermal delivery

## Abstract

Solid microneedles can successfully puncture the stratum corneum and thus enable the drugs to migrate from the adhesive patch to the viable skin tissues for therapy. The treatment in different skin layers can vary greatly. However, how to improve its effectiveness remains less understood. In this study, numerical simulation is employed to predict the transport and disposition of drugs in each skin layer and blood using a skin model rebuilt from the real skin anatomical structure. The therapeutic effect is assessed by exposure to drugs over time. Results reveal the dominance of diffusion in determining the transport of nanosized drug carriers and free drugs in viable skin tissues. Delivery outcomes are highly sensitive to drug delivery system properties. Increasing the nanocarrier partition coefficient or diffusion coefficient in the skin can successfully enhance the treatment in entire skin tissue and blood. The enhancement can also be obtained by reducing the microneedle spacing or patch thickness. However, several properties should be optimised individually with respect to the target site’s location, including the microneedle length, diffusion coefficient of nanocarriers in the skin, drug release rate and nanocarrier vascular permeability. Drug concentrations in the blood can be effectively increased when administered to skin areas rich in capillaries; whereas, the treatment in the skin tissues slightly would reduce simultaneously. Furthermore, delivery results are insensitive to changes in lymphatic function and the properties of free drugs introduced by the medicated patch. These findings can be used to improve transdermal drug delivery for better treatment.

## 1. Introduction

Transdermal delivery is a potential alternative to hypodermic injection and oral delivery to treat a variety of diseases including diabetes [1] and Alzheimer’s disease [2]. In conventional transdermal delivery where administration happens on the skin surface, drugs need to pass through the stratum corneum (SC) and viable epidermis (VE) to arrive at the papillary dermis (PD), in which capillaries are mainly embedded. These blood vessels allow the drugs to enter the circulatory system and then become available for systemic absorption, while the remaining drugs in the papillary dermis can travel further down to the deeper skin tissue of the reticular dermis (RD). Transdermal delivery is favoured due to its advantage in providing continuous drug administration without repeated invasive procedures [3]. However, its performance is greatly limited by the stratum corneum which is almost impenetrable to most substances [4]. Solid microneedles are an effective transdermal drug delivery system that can overcome this obstacle. In the treatment, the stratum corneum is punctured by solid microneedles first and a medicated adhesive patch (PT) is then imposed on the skin surface after the microneedles have been withdrawn. The loaded drugs can travel from the patch to the cavities (CV) which are left by the microneedles, and thence enter the viable skin tissue. Various types of solid microneedles are available to meet different treatment requirements [5,6]. However, the treatment still needs to be improved since metabolic reactions in skin tissues can quickly eliminate the administrated drugs and thereby reduce drug penetration depth and deposition, particularly in deep skin tissues. Nanocarriers are developed to encapsulate drugs inside using materials that are inert to undesired bioreactions. By slowly releasing the payloads, these drug vehicles can effectively slow down the elimination of small-molecule drugs, enabling a long-lasting drug supply [7]. Solid microneedles have been combined with nanocarriers in preclinical trial studies on transdermal drug delivery [8], but the impact of multiple factors that can affect the delivery results has not been determined.

Transdermal drug delivery involves a series of interrelated physiological and physicochemical steps, making it less feasible to examine each influencing factor by clinical means. Numerical simulation provides a solution. Using a customised mathematical model to describe the specific drug transport steps, the simulation enables conducting comprehensive parametric studies to identify the impact of each factor in an individual or integrated manner and optimise the drug delivery systems [9]. This research method plays an increasingly important role in the research of transdermal drug delivery. It was employed to simulate drug transport from the viable epidermis to the reticular dermis to identify the influences of the size, depth, blood flow rate, and density of capillaries in the papillary dermis [10]. An application of simulating transdermal delivery using dissolving microneedles was given in Ref. [11], where the predicted drug concentrations well agreed with the experimental data. Transdermal delivery using drug-loaded microneedles was numerically investigated under different conditions to reveal the influences of different drug delivery system properties and environments [12]. However, simulations of transdermal delivery using solid microneedles are less reported.

In this study, solid microneedle-mediated transdermal drug delivery is studied under various delivery conditions using a mathematical model covering the critical transport processes; these include the flow of interstitial fluid, exchange of fluid and drug between the different skin layers and circulatory systems, drug release from nanocarriers, convective and diffusive drug transport in association the and tissues, drug-protein disassociation, drug elimination owing to metabolic reactions, drug plasma clearance and physical degradation. Treatment in every skin layer and the blood is assessed by the parameter drug exposure over time which is calculated based on the local drug concentration predicted from the model.

## 2. Materials and methods

### 2.1 Mathematical model

Stratum corneum contains multiple corneocyte layers that are sealed by lipid matrixes, making it impenetrable to a variety of substances including most therapeutic compounds and agents. Although water molecules can cross the stratum corneum to the environment, this transport termed trans-epidermis water loss highly depends on the difference between the saturated vapour pressure of water at the skin surface and ambient relative humidity and is often treated differently from interstitial fluid flow [13]. Since it is covered by a medicated adhesive patch all the time in the solid microneedle-mediated transdermal delivery, the stratum corneum is assumed to be saturated. A mathematical model was established in our previous study on the delivery using drug-loaded microneedles [12]. This model is further developed to accommodate the specific drug transport steps induced by the combined use of solid microneedles and medicated adhesive patches. The governing equations are summarised below.

#### 2.1.1 Model of interstitial fluid flow

The cavities left after the microneedles are withdrawn are filled with interstitial fluid. Viable skin tissues can be treated as porous media in which interstitial fluid flows in the gaps between cells. So that the transport of Newtonian, incompressible interstitial fluid at the quasi-steady state is modelled by the mass and momentum equation, as

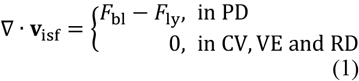

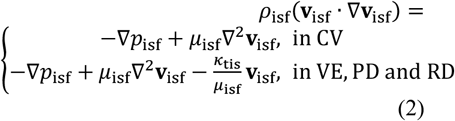

in which *ρ*_isf_ and *μ*_isf_ are the interstitial fluid density and viscosity, respectively. *p*_isf_ refers to the interstitial fluid pressure, and **v**_isf_ is the flow velocity. *κ*_tis_ stands for the permeability of skin tissues. Capillaries which are homogeneously distributed in the papillary dermis are considered a source term. The fluid exchange between the papillary dermis and blood (*F*_bl_) can be described by Starling’s law in the form of

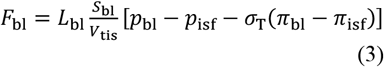

in which *L*_bl_ and *S*_bl_ are the hydraulic conductivity and surface area of blood capillary walls, respectively. *V*_tis_ is the local tissue volume. The blood pressure is represented by *p*_bl_. *σ*_T_ is the osmotic reflection coefficient. *π*_isf_ and *π*_bl_ are the osmotic pressure of interstitial fluid and blood, respectively. Lymphatic vessels are present in the papillary dermis, running parallel to the blood vessels [9]. The fluid loss from the skin to the lymphatic system (*F*_ly_) can be expressed by

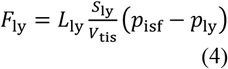

where *L*_ly_ and *S*_ly_/*V*_tis_ have the same definitions as those of blood vessels. *p*_ly_ stands for the pressure of lymph.

#### 2.1.2 Drug transport model

The transport of nanocarriers and released drugs in the patch, microneedle-induced cavity, skin tissues, and circulatory systems are schematically depicted in **Figure 1**. The letters NC, FD and BD stand for nanocarriers, free drugs and protein-bound drugs, respectively.

**Figure 1.**
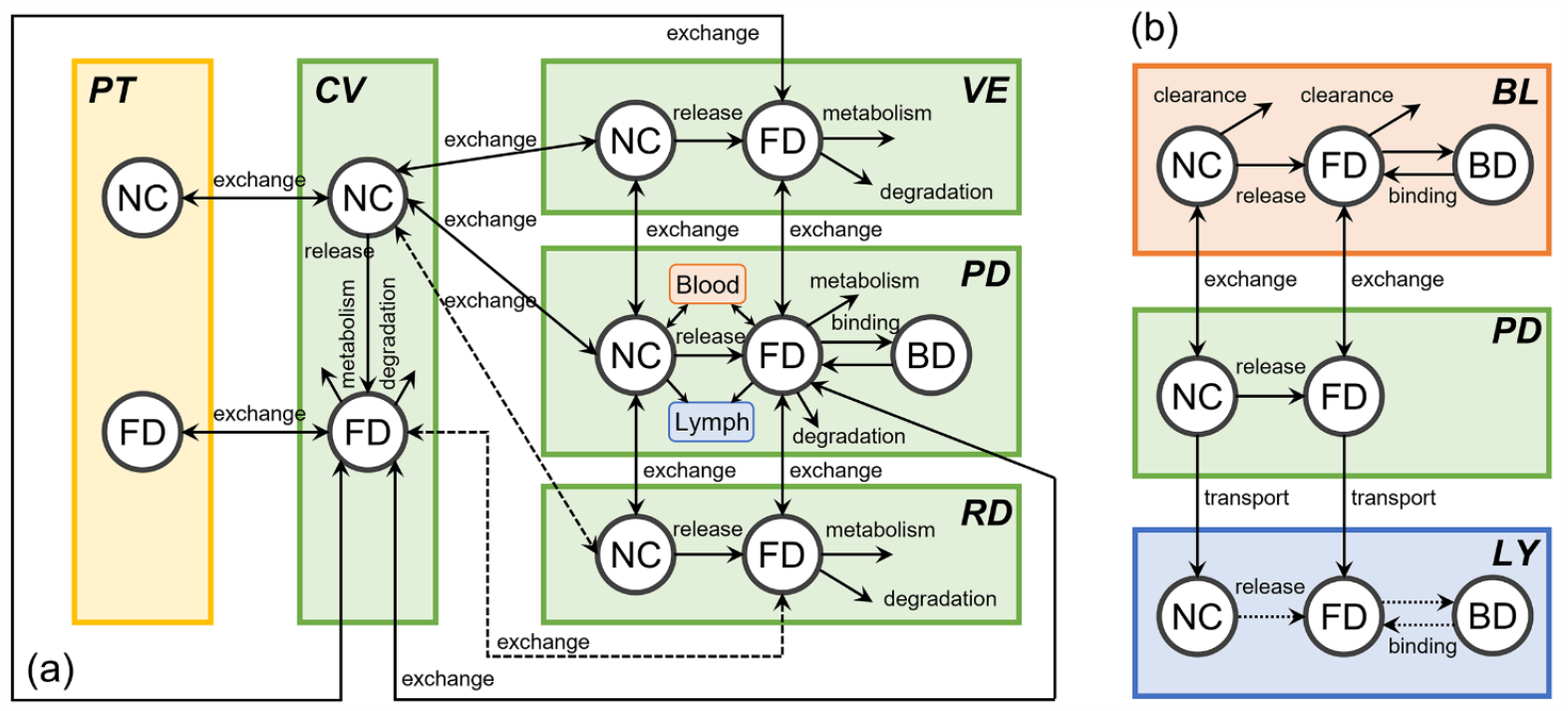
Drug transport in the transdermal delivery of drug nanocarriers using solid microneedles. (a) the overarching depiction of the key transport steps among the patch, skin and circulatory systems. (b) a close look at drug transport between the skin tissue, blood and lymphatic circulatory systems. The lymphatics is considered to be a sink. Hence, the drug release dynamics and drug binding protein in the lymph are not specified in the simulation, shown as the dotted lines. Drug transport between the cavity and reticular dermis only occurs if the solid microneedles are able to reach this skin layer. Therefore, these two drug transport processes are shown as the dashed lines. The two interactions of drug binding and unbinding with proteins are only considered in the circulatory systems and papillary dermis, since the proteins, e.g. albumin, are mainly transported by the blood, and blood capillaries are embedded in this skin layer. This diagram schematically shows the key drug transport processes, not presenting the real thickness and depth of the skin layers.

##### 2.1.2.1 Drug transport in patch

Small-molecule drugs are considered well-encapsulated within nanocarriers before travelling to the cavity. The concentration of nanocarriers in a patch (*C*_NC,PT_) is determined by diffusion, as

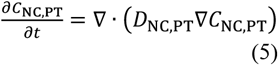

where *D*_NC,PT_ is the diffusion coefficient of nanocarriers in the patch. *t* is time. Free drugs can travel back to the patch after being released into the skin. The free drug concentration in the patch (*C*_FD,PT_) can be calculated by

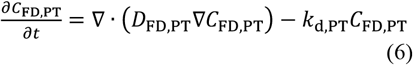

where *k*_d,PT_ is the rate of physical degradation in the patch. *D*_FD,PT_ is the local diffusion coefficient of free drugs.

##### 2.1.2.2 Drug transport in cavity

The travel of nanocarriers in the microneedle-induced cavity is subject to concentration gradient-driven diffusion and convection with the flow of interstitial fluid. The concentration (*C*_NC,CV_), which also depends on drug release, can be calculated by

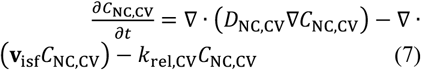

in which *D*_NC,CV_ and *k*_rel,CV_ are the nanocarrier diffusion coefficient and release rate in the cavity, respectively. The movement of free drugs in the cavity (*C*_FD,CV_) is subject to diffusion and convection, physical degradation, metabolic reactions and drug release, as

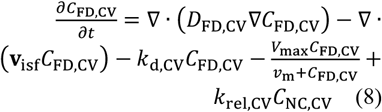

where *D*_FD,CV_ refers to the diffusion coefficient of free drugs in the cavity. *V*_max_ and *ν*_m_ are the metabolic reaction constants. *k*_d,CV_ is the free drug degradation rate in the cavity.

##### 2.1.2.3 Drug transport in viable epidermis

The concentration of nanocarriers in the viable epidermis (*C*_NC,VE_) is determined by local drug release and drug transport by convection and diffusion, as

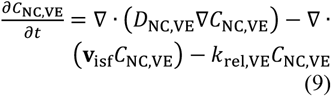

in which *D*_NC,VE_ and *k*_rel,VE_ are the diffusion coefficient of nanocarriers and drug release rate in the viable epidermis, respectively. The free drug concentration (*C*_FD,VE_) in the viable epidermis depends on convection, diffusion, physical degradation, metabolic reactions and local drug release, as

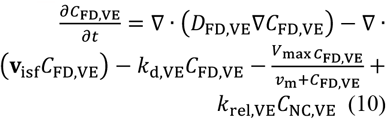

where *D*_FD,VE_ is the diffusion coefficient of free drugs in the viable epidermis.

##### 2.1.2.4 Drug transport in papillary dermis

Nanocarriers move through the papillary dermis via diffusion and convection. Since the drug loss to the blood and lymphatic systems, and local drug release also contribute to the nanocarrier deposition in this layer, the concentration (*C*_NC,PD_) can be obtained by

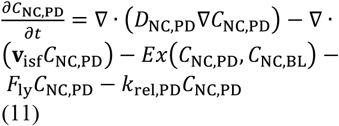

where *D*_NC,PD_ and *k*_rel,PD_ are the nanocarrier diffusion coefficient and release rate in the papillary dermis, respectively. *Ex*(*C*_NC,BL_, *C*_NC,PD_) is the rate of nanocarrier exchange between the papillary dermis and blood, defined as

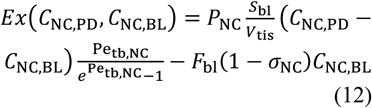

where *P*_NC_ is the nanocarrier’s effective vascular permeability. *σ*_NC_ is the reflection coefficient of nanocarriers. *C*_NC,BL_ is the blood concentration of nanocarriers. Pe_tb,NC_ is the Péclet number of nanocarriers when passing through the capillary walls, as

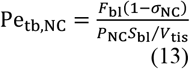

The free drug concentration (*C*_FD,PD_) in the papillary dermis is subject to diffusive and convective drug transport, drug physical degradation, metabolic reactions, drug exchange between the papillary dermis and blood, drug loss to the lymphatic system, binding with proteins and drug release, as

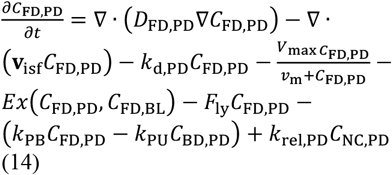

where *D*_FD,PD_ is the diffusion coefficient of free drugs in this layer. *k*_PB_ and *k*_PU_ are the protein-drug binding and unbinding rates, respectively. *k*_d,PD_ stands for the local drug degradation rate. The free drug exchange rate, *Ex*(*C*_FD,PD_, *C*_FD,BL_) is defined the same as in **Equation (12)** using the properties and concentration of free drugs. The blood concentration of free drugs is *C*_FD,BL_ . The bound drug concentration in this layer (*C*_BD,PD_) can be calculated using

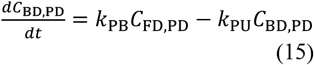

##### 2.1.2.5 Drug transport in reticular dermis

Nanocarriers transfer by diffusion and convection in the reticular dermis. Its concentration (*C*_NC,RD_) also depends on drug release from nanocarriers, as

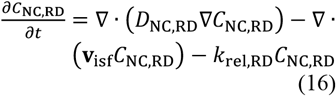

where *D*_NC,RD_ and *k*_rel,RD_ are the local nanocarrier diffusion coefficient and drug release rate, respectively. The free drug concentration in the reticular dermis (*C*_FD,RD_) is subject to diffusion and convection, physical degradation, metabolic reactions and drug release, as

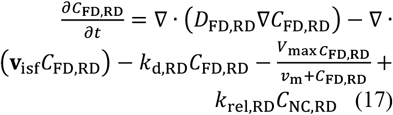

where *k*_d,RD_ is the local drug degradation rate. *D*_FD,RD_ is the diffusion coefficient of free drugs.

##### 2.1.2.6 Drug transport in blood

Nanocarriers can continue releasing the loaded drugs after entering the blood circulation system and are eliminated by kidneys and other organs. The blood concentration of nanocarriers (*C*_NC,BL_) can be obtained by

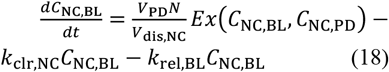

in which *V*_PD_ is the papillary dermis volume that is covered by each solid microneedle. *V*_dis,NC_ stands for the nanocarrier distribution volume. *N* refers to the total number of solid microneedles in an array. *k*_rel,BL_ is the drug release rate in the blood. *k*_clr,NC_ is the clearance rate of nanocarriers in blood.

The blood concentration of free drugs (*C*_FD,BL_) is subject to the drug exchange between the blood and papillary dermis tissue, binding and unbinding with proteins, plasma clearance and drug release, as

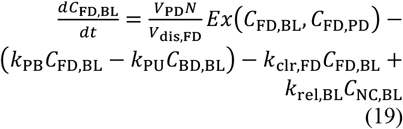

where *k*_clr,FD_ and *V*_dis,FD_ are the rate of plasma clearance and distribution volume of free drugs, respectively. The blood concentration of protein-bound drugs (*C*_BD,BL_) can be calculated by

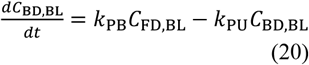

### 2.2 Model geometry

Because multiple solid microneedles are usually arranged in an array at the same distance, a representative elementary volume (REV) is chosen for numerical simulations, as shown in **Figure 2**. The 2D axis-symmetric configuration comprises a patch that loads nanocarriers, a cavity left after the withdrawal of the microneedle, and the skin layers; their real thicknesses [10] are specified in **Figure 2**. The morphological features of solid microneedles can vary significantly depending on a particular design. In this study, 10 × 10 tapered solid microneedles with a length of 400 μm are spaced 600 μm apart in the array. The radius at the microneedle base is 150 μm. This microneedle is long enough to pierce the superficial layers and reach the papillary dermis. The thickness of the medicated adhesive patch is 100 μm [8]. There are approximately 70,000 triangular elements in the computational mesh based on the mesh sensitivity study. The finest elements which have a size of 0.005 μm are applied at the patch-cavity and cavity-tissue interfaces.

**Figure 2.**
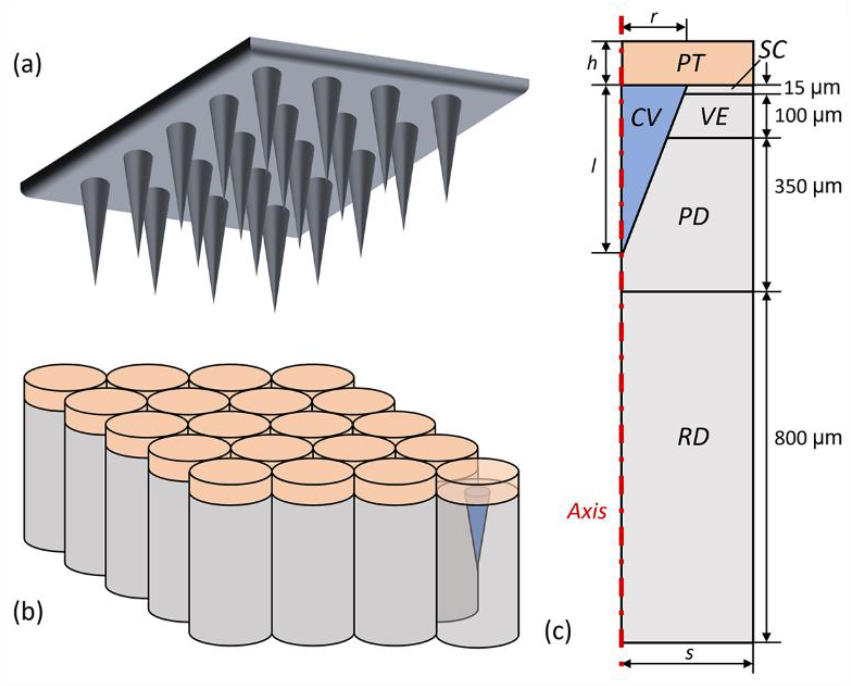
Model geometry. Schematical diagram of solid microneedle array (a), the array of representative elementary volumes and (c) the computation domain. The skin tissues, cavity and medicated adhesive patch are in light grey, blue and pink, respectively. *l* is the microneedle length. *r* is the radius of the microneedle base. *h* is the patch thickness. *s* is the radius of representative elementary volume, which is half of the tip-to-tip distance.

### 2.3 Model parameters

The drug administration duration is usually much shorter compared to the time window of tissue growth. Therefore, the tissue properties and drug properties are considered constants which do not change with time. Doxorubicin is chosen as the representative drug. The base values of tissue and drug properties are given in **Table 1** and **Table 2**, respectively. The ranges of each key property are specified in the following sections where their roles are studied.

**Table 1.** Properties of skin tissues

| Symbol | Parameter | Unit | VE | PD | RD | Source |
| --- | --- | --- | --- | --- | --- | --- |
| $\kappa_{tis}$ | Tissue hydraulic permeability | $m^2$ | $1.0 \times 10^{-16}$ | $1.0 \times 10^{-16}$ | $1.0 \times 10^{-16}$ | [15] |
| $\mu_{is}$ | Interstitial fluid viscosity | $Pa \cdot s$ | $7.8 \times 10^{-4}$ | $7.8 \times 10^{-4}$ | $7.8 \times 10^{-4}$ | [38] |
| $\rho_{is}$ | Interstitial fluid density | $kg/m^3$ | $1.0 \times 10^3$ | $1.0 \times 10^3$ | $1.0 \times 10^3$ | [39] |
| $\pi_{bl}$ | Blood osmotic pressure | $Pa$ | — | $2.7 \times 10^3$ | — | [40] |
| $\pi_{is}$ | Osmotic pressure of interstitial fluid | $Pa$ | — | $1.3 \times 10^3$ | — | [40] |
| $\sigma_T$ | Osmotic reflection coefficient | — | — | $9.1 \times 10^{-1}$ | — | [40] |
| $L_{bl}$ | Hydraulic conductivity of capillary walls | $m/Pa/s$ | — | $2.7 \times 10^{-12}$ | — | [40] |
| $p_{bl}$ | Pressure in capillary | $Pa$ | — | $2.1 \times 10^3$ | — | [40] |
| $S_{bl}/V_{tis}$ | Surface area of capillary in unit tissue volume | $m^{-1}$ | — | $6.0 \times 10^3$ | — | [32] |
| $L_{ly}S_{ly}/V_{tis}$ | Transport rate of interstitial fluid to lymphatic system | $Pa^{-1}s^{-1}$ | — | $4.2 \times 10^{-7}$ | — | [40] |
| $p_{ly}$ | Pressure in lymphatic system | $Pa$ | — | 0 | — | [40] |

**Table 2.** Properties of nanocarriers and free drugs

| Symbol | Parameter | Unit | Nanocarrier | Free drug |
| --- | --- | --- | --- | --- |
| $D_{PT}$ | Diffusion coefficient in cavity | $m^2/s$ | $2.9 \times 10^{-12}$ [29] | $3.8 \times 10^{-10}$ [41] |
| $D_{tis}$ | Diffusion coefficient in PT, VE, PD and RD | $m^2/s$ | $1.0 \times 10^{-13}$ [15] | $1.0 \times 10^{-10}$ [15] |
| $K$ | Partition coefficient between patch and cavity | — | 1.0 [9] | 1.0 [9] |
| $k_{rel}$ | Drug release rate | $s^{-1}$ | $1.0 \times 10^{-4}$ [42] | — |
| $k_d$ | Physical degradation rate of free drug | $s^{-1}$ | — | $5.6 \times 10^{-6}$ [43] |
| $V_{max}$ | Michaelis–Menten constant for metabolic reaction | $mol /m^3/s$ | — | $5.1 \times 10^{-1}$ [44] |
| $v_m$ | Michaelis–Menten constant for metabolic reaction | $mol/m^3$ | — | $6.7 \times 10^{-3}$ [44] |
| $\sigma$ | Osmotic reflection coefficient | — | 1.0 [45] | $1.5 \times 10^{-1}$ [38] |
| $k_{PB}$ | Binding rate of drugs with proteins | $s^{-1}$ | — | $8.3 \times 10^{-1}$ [46] |
| $k_{PU}$ | Unbinding rate of drugs with proteins | $s^{-1}$ | — | $2.8 \times 10^{-1}$ [46] |
| $P$ | Effective vascular permeability | $m/s$ | $1.0 \times 10^{-9}$ [25] | $3.8 \times 10^{-7}$ [15] |
| $C_{in}$ | Administrated dose | $M$ | 1.0 [47] | — |
| $k_{clr}$ | Plasma clearance rate | $s^{-1}$ | $5.0 \times 10^{-5}$ [15] | $1.0 \times 10^{-4}$ [15] |
| $V_{dis}$ | Distribution volume | $m^3$ | $1.8 \times 10^{-2}$ [15] | $2.0 \times 10^{-2}$ [15] |

### 2.4 Boundary conditions

Owing to the stratum corneum’s nearly impenetrable nature, the drug fluxes are assumed to be zero at the boundaries between the stratum corneum and its adjacent compartments, including the patch, cavity and viable epidermis, as

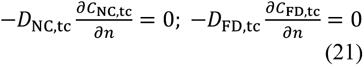

where the subscript tc stands for the adjacent compartments depending on the location. *n* is the normal direction of the boundary. The top surface of the patch is assumed to be impermeable to the drugs and water molecules to maintain the loaded drugs within the patch. The travel of therapeutic agents across the interface between the patch and cavity is governed by

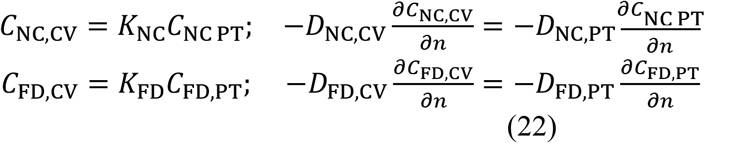

in which *K* refers to the partition coefficient of therapeutic agents between the cavity and patch. Given viable skin tissues are aqueous phases [9], variables at the boundaries among the cavity and different skin layers are considered to be continuous [14,15]. The symmetric boundary condition is imposed on the REV side. The fluxes of interstitial fluid and drugs are zero at the bottom of the domain [10].

### 2.5 Numerical methods

The governing equations in the mathematical model are solved using COMOSL multiphysics (COMSOL Inc., Stockholm, Sweden). The time step is set to be 0.1 ms based on the time-step sensitivity study. The interstitial fluid flow model is first applied to predict the flow pressure and velocity, which are then sent to the drug transport model to simulate drug penetration and deposition. All drugs are within the nanocarriers homogeneously distributed in the patch at the beginning of the transdermal delivery.

### 2.6 Quantification of delivery results

#### 2.6.1 Spatially averaged concentration

The drug deposition in a specific compartment including each skin layer and blood can be assessed by the spatially averaged concentration (*C*_avg_), defined as

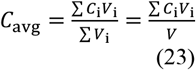

in which the subscript i indicates the local variable. *V* is the volume of the entire studied compartment.

#### 2.6.2 Exposure to drugs over time

The treatment effectiveness in each compartment can be presented by the exposure to drugs over time, *AUC*, which is calculated as the integration of free drug concentration over time, as

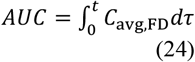

## 3. Results

### 3.1 Baseline delivery

The nanocarriers and released drugs move in the interstitial fluid after escaping from the patch. The predicted flow velocity in the cavity and skin tissues is on the order of 10^−8^ nm/s and 10^−13^ nm/s, respectively. This significantly low velocity can be attributed to the impermeable patch which prevents transepidermal water loss from the skin surface to the environment. The Péclet number, Pe = *ν*_isf_*L*_c_/*D*, can be used to evaluate the importance of convection and diffusion in determining drug transport. *ν*_isf_ and *L*_c_ are the scale of interstitial fluid velocity and transport length, respectively, and *D* is the diffusion coefficient. It is calculated as 3.5 × 10^−10^ and 2.6 × 10^−12^ for nanocarriers and free drugs, respectively, in the cavity, and 1.0 × 10^−13^ and 1.0 × 10^−16^ in the skin tissues. Since all the values are much lower than 1.0, this calculation denotes that drug transport is dominated by diffusion in the transdermal drug delivery using solid microneedles.

**Figure 3**(a) shows the spatial distribution of nanocarriers in the patch and skin at different time points. Loss of nanocarriers from the patch starts at the patch-cavity interface and continuously spreads to the patch margin, resulting in a sharp concentration gradient in the patch at the beginning of the treatment. With time proceeding, this concentration distribution becomes more uniform owing to the diffusion of nanocarriers. As shown in **Figure 3**(b), although the concentration in the patch is much higher, free drugs can successfully disperse in the cavity and travel to the surrounding skin tissues.

**Figure 3.**
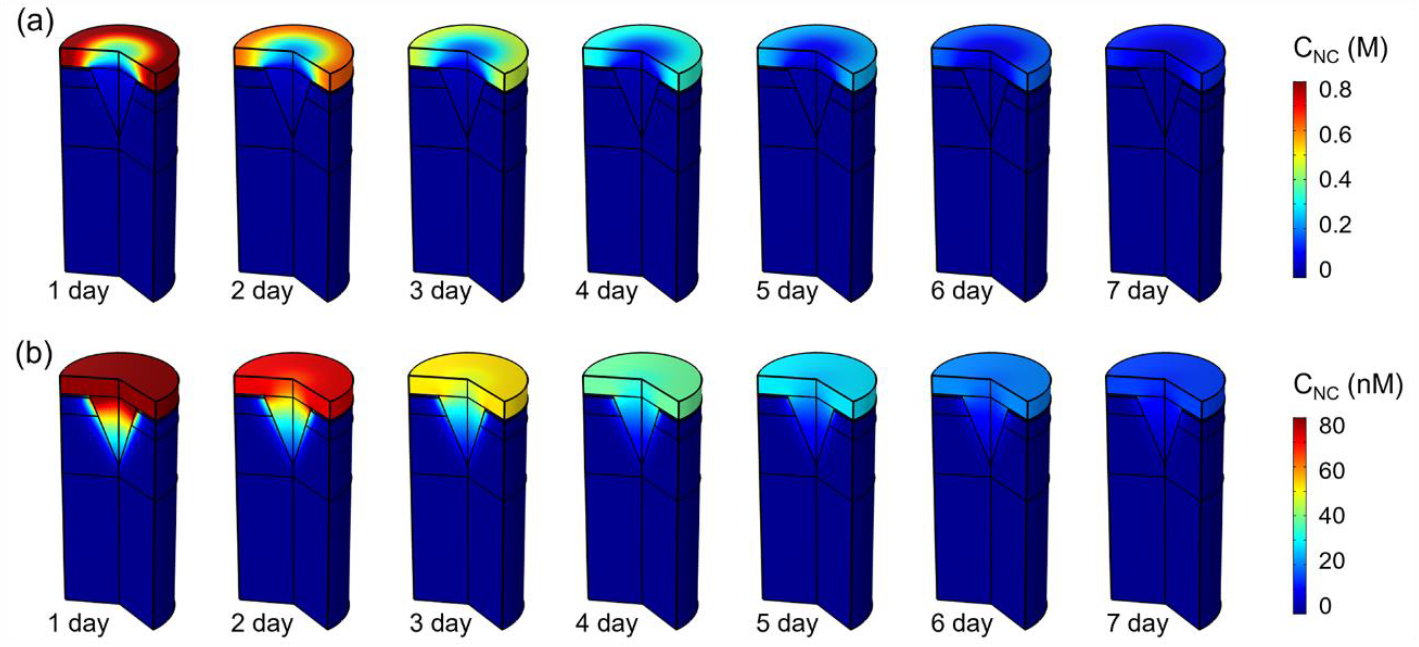
Spatial distribution of therapeutic agents in the patch, cavity and skin tissues at different time points. (a) nanocarriers and (b) free drugs.

**Figure 4** displays the spatially averaged concentrations of different forms of drugs in the patch, cavity, skin layers and blood as a function of time. The patch concentration of nanocarriers decreases exponentially over time due to the continuous loss from the patch to the skin. As a result, the nanocarrier concentration in the cavity quickly rises to the peak in about 2.8 hours and then gradually decreases as time proceeds. This concentration presents a similar trend in the viable epidermis as well as papillary dermis. In contrast, the concentration in the reticular dermis and blood remains low over time, indicating that few nanocarriers can travel to the deep tissues for therapy. Moreover, the accumulation of free drugs shows similar trends in all compartments, with the order of concentration from high to low consistent with their depth in the skin. Notably, the free drug concentration is 1/3 of that of bound drugs all the time, determined by the drug binding kinetics and drug properties in **Table 2**. So the analysis of the next simulations will focus on nanocarriers and free drugs.

**Figure 4.**
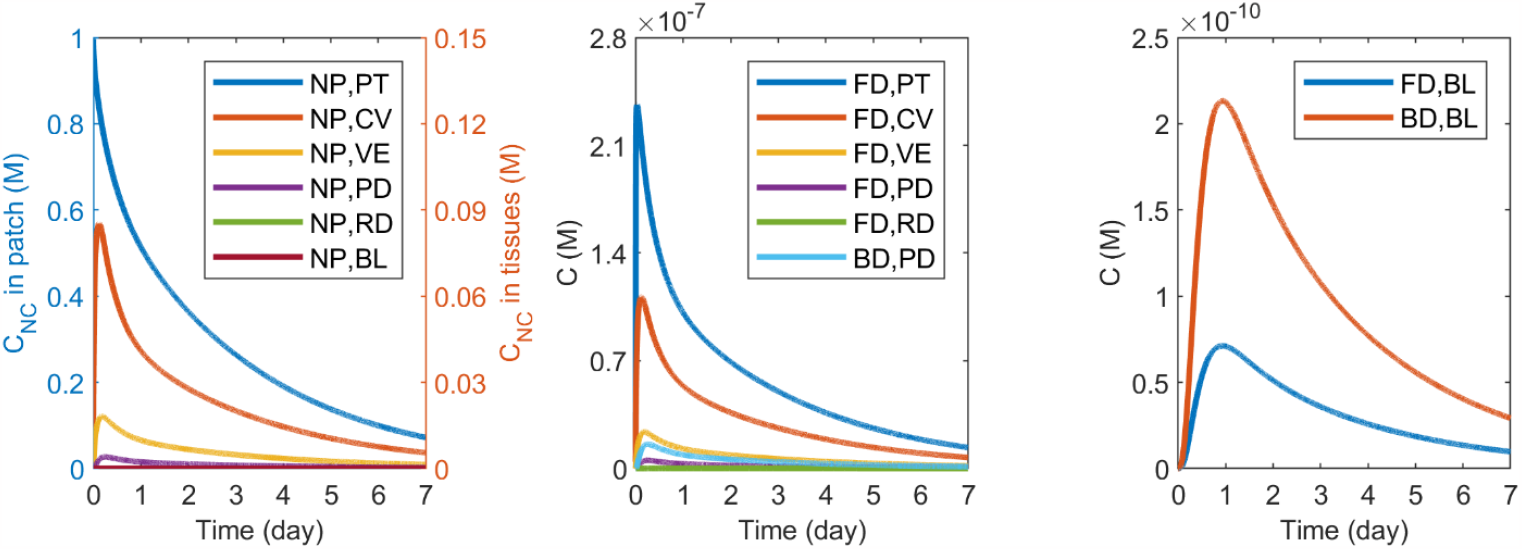
Time courses of spatially averaged concentrations of drugs in different forms. (a) the nanocarrier concentrations in each compartment of the computational domain. The concentration of free drugs and drugs that are bound with proteins in the patch and skin (b) and blood (c).

### 3.2 Role of microneedle properties

#### 3.2.1 Microneedle length

Microneedle length directly defines the penetration depth in the skin and hence delivery results. The selection of this morphological feature requires the consideration of multiple factors, such as the location of the target site and skin tissue thickness. The microneedles are usually hundreds of micrometres long [6,16]. In this study, this parameter changes from 100 to 700 μm to examine its impact. Specifically, the microneedle with a length of 100 μm only stays at the viable epidermis; whereas, the 250 μm and 400 μm microneedles can reach the papillary dermis. Penetration into the reticular dermis can be obtained using the 550 μm and 700 μm microneedles. Please note that the microneedle radius is changed simultaneously with its length to keep the volume of microneedles and cavities identical. **Figure 5** shows how microneedle length influences the delivery to different skin layers and the blood. The loss of nanocarriers in the patch can be effectively slowed down by elongating the microneedle because the shrinkage of the microneedle radius leads to a decrease in the ratio of patch-cavity interface area to patch volume. As a result of this slow but durable drug supply from the patch, the concentration in the cavity and viable epidermis rise to lower peaks but decrease more gradually over time. The peak concentration in the papillary dermis is nonlinearly related to the microneedle length, with the highest occurring when the microneedle stays at this particular skin layer. The concentration in the blood shows a similar trend since the capillaries are mainly present in the papillary layer. The reticular dermis is originally the most difficult to receive drugs. Using the microneedles that can reach this layer is able to significantly improve the local drug accumulation. Comparisons on drug exposure demonstrate that the treatment for viable epidermis decreases with the length of microneedles. The 400 μm microneedle is the optimum to improve drug delivery to the papillary dermis and blood circulatory system. However, treatment in the reticular dermis responds positively to the elongation of microneedles; *AUC* in the reticular dermis rises sharply when the microneedle is capable of piercing the superficial tissues to reach this layer.

**Figure 5.**
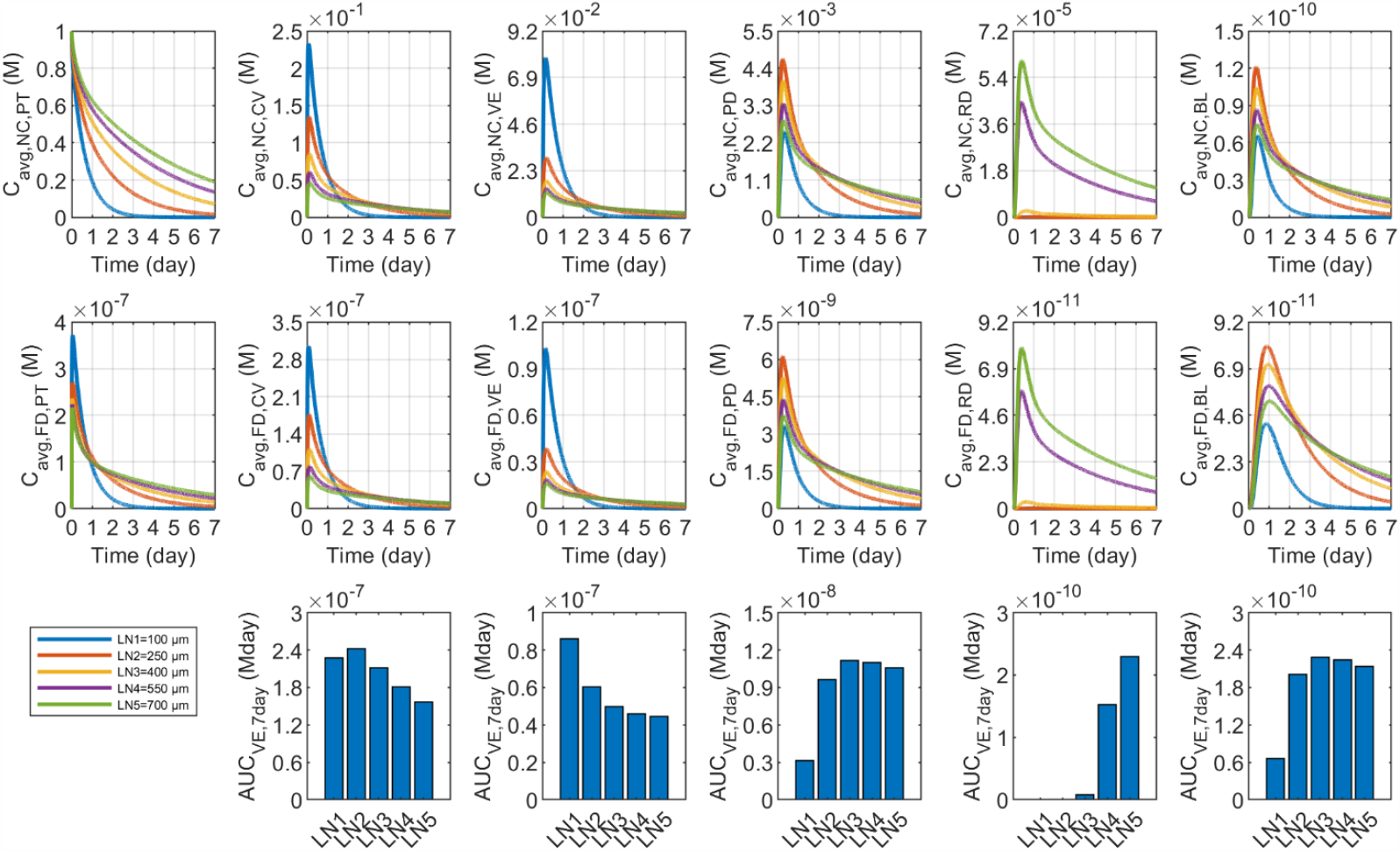
Impact of microneedle length (*l*) on the transdermal delivery results using solid microneedles. Upper panel: spatially averaged concentration of nanocarriers; Middle panel: spatially averaged concentration of free drugs; Lower panel: drug exposure over time. The columns from left to right in turn are for the patch (PT), cavity (CV), viable epidermis (VE), papillary dermis (PD), reticular dermis (RD) and blood (BL). The same plot playout is applied to the following figures where the impact of each property is discussed.

#### 3.2.2 Microneedle spacing

The distance between two microneedles is another parameter besides microneedle length that can be tailored in fabrication. This parameter was 600 μm in Ref.[17], while the microneedles were 1000 μm away from each other in Ref.[18]. Therefore, this microneedle property is varied in the range from 400 μm to 1200 μm to determine its role. Please note that the initial patch concentration of nanocarriers is changed simultaneously with microneedle spacing to keep the dosing identical.

The results in **Figure 6** show that the spatially averaged concentration of nanocarriers in the patch drops faster as the distance between microneedles decreases. This can be attributed to the reduction in the patch volume in REV as a result of the shortening of the microneedle spacing. On the one hand, the increased nanocarrier concentration leads to a sharper concentration gradient across the cavity-patch interface, accelerating the nanocarrier loss from the patch; On the other hand, since the size of the microneedles remains constant, the ratio of patch-cavity interface area over the patch volume becomes higher as the microneedles get closer, resulting in more efficient transport of nanocarriers from the patch to the skin. Consequently, when microneedles are brought closer together, the nanocarrier concentration in all downstream compartments can increase more rapidly to higher peaks, but also rapidly decrease to lower levels over time. This is primarily because of the high initial nanocarrier concentration in the patch and the rapid but unsustainable nanocarrier loss to the skin. The free drug concentration in each skin layer and blood exhibit similar trends because all free drugs are released from nanocarriers. Drug exposure depends on the peak the free drug concentration can reach and the change rate of concentration over time. The lower panel shows that treatment in the whole skin tissue and blood decreases with increasing microneedle spacing when the dose remains constant.

**Figure 6.**
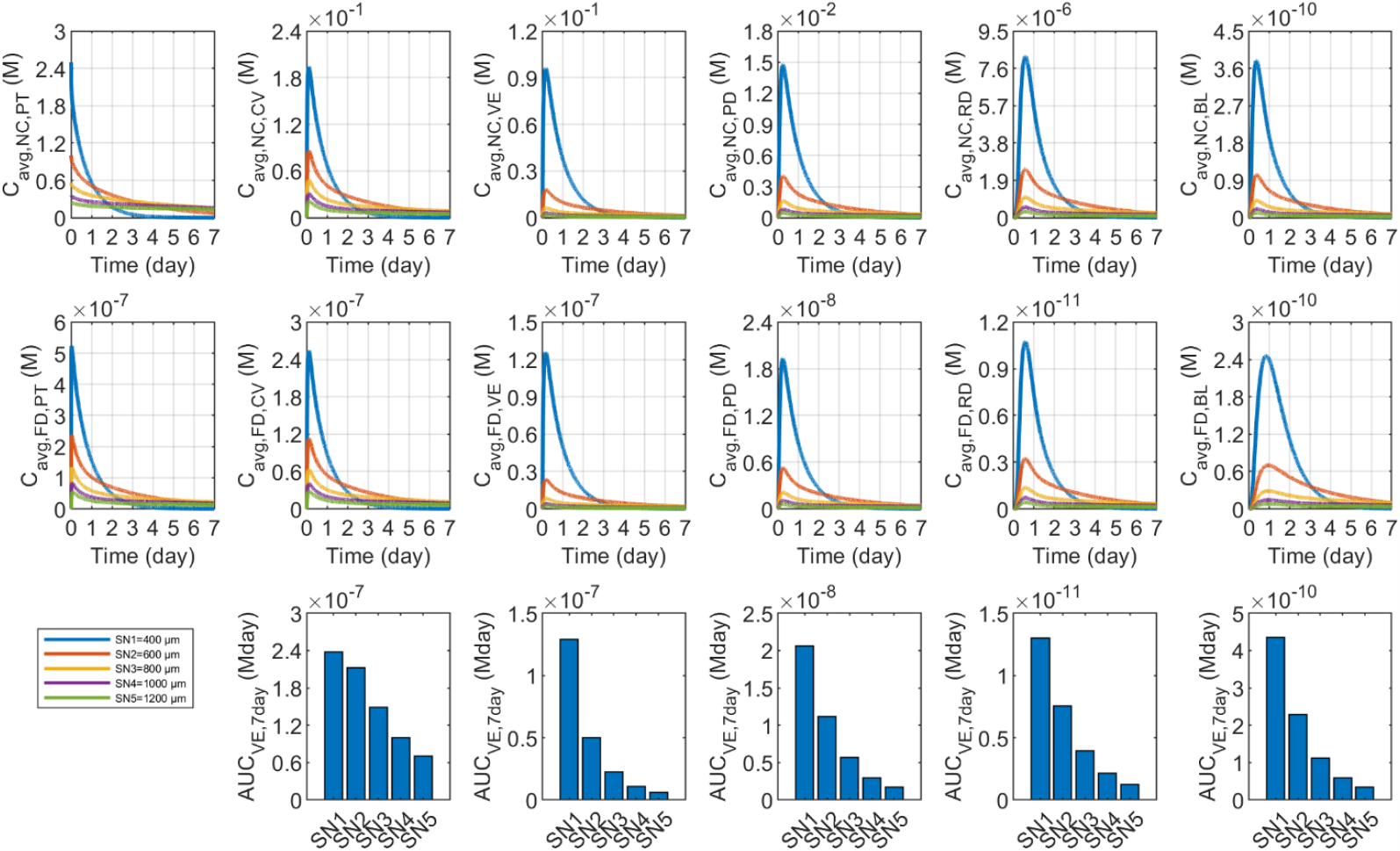
Impact of microneedle spacing (*s*) on the transdermal delivery results using solid microneedles.

### 3.3 Role of nanocarrier properties

#### 3.3.1 Drug release rate

The rate of drug release from nanocarriers is a crucial factor in controlled drug delivery systems. It represents the time scale in which the drug vehicles release the payloads, strongly determining the therapeutic effects. Its value can vary considerably depending on the nanocarrier formulation and surrounding environment. The first-order kinetics is usually used to derive this parameter from the cumulative drug release profiles [19]. For instance, this parameter was calculated as 9.5 × 10^−5^ s^−1^ for the pH-sensitive nanocarriers in Ref.[20]. Thermosensitive liposomes can release the loaded drugs at the rate of 5.4 × 10^−2^ s^−1^ once the temperature is greater than the phase transition threshold of the lipid membrane [21]. Therefore, the drug release rate is varied in a wide range from 1.0 × 10^−6^ s^−1^ to 1.0 × 10^−1^ s^−1^ in this study.

**Figure 7** shows how the drug release rate influences the delivery results. It is not surprising that nanocarrier concentration in every compartment reduces when drug release is accelerated. A burst of free drug concentration occurs in the viable epidermis at the onset of treatment when the release rate is high. However, this concentration would drop rapidly as a result of drug elimination and drug transport into the surrounding skin tissues. A similar trend can be found in the patch due to drug exchange between these two adjacent compartments of the patch and cavity. The response of free drug concentration to the changes in release rate differs distinctly between the skin layers. On the one hand, the rapid release results in a large amount of drugs presenting in their free form in the skin, especially the superficial layers, and thence enhances the diffusion of free drugs into deep tissues. On the other hand, fewer nanocarriers can travel to the deep tissues to release the payload. The treatment efficacy in each layer, evaluated in terms of *AUC*, varies the drug release rate. Raising the drug release rate to 1.0 × 10^−4^ s^−1^ enables more effective treatment in the viable epidermis, whereas the optimal rate is found to be 1.0 × 10^−3^ s^−1^ for the rest skin layers and systemic absorption via blood.

**Figure 7.**
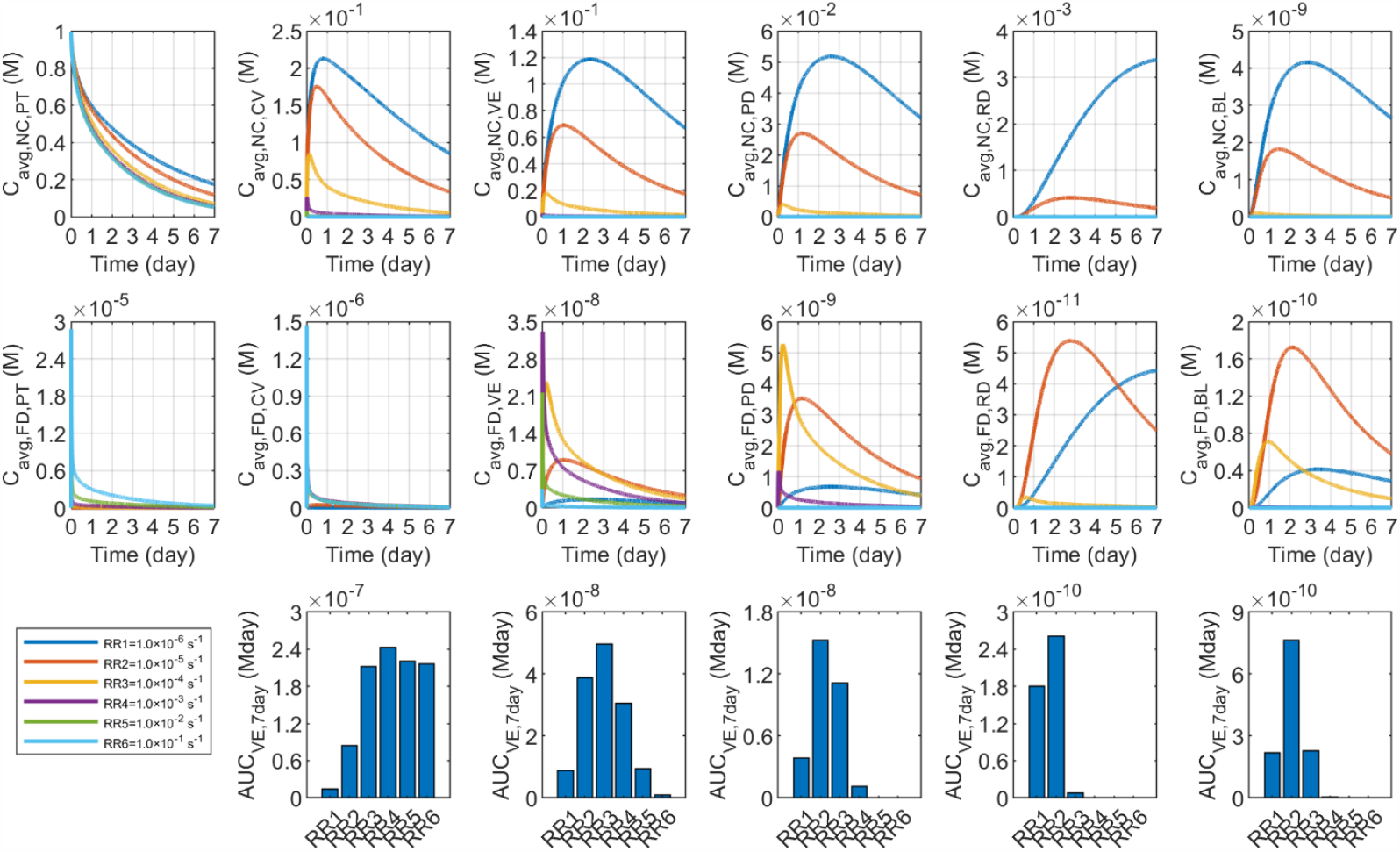
Impact of drug release rate (*k*_rel_) on the transdermal delivery results using solid microneedles.

#### 3.3.2 Diffusion coefficient of nanocarriers in skin

This diffusion coefficient describes the nanocarriers’ transport ability in tissues by thermal motion, greatly dependent on their dimensions. This parameter was reported to be 2.4 × 10^−13^ m^2^/s [22] and 1.0 × 10^−14^ m^2^/s [23] for the 100 nm and 500 nm nanocarriers, respectively. The diffusion coefficient of a 3 nm nanocarrier was found to be 2.2 × 10^−11^ m^2^/s in the experiment using agarose phantom [24]. Therefore, a large range from 1.0 × 10^−15^ to 1.0 × 10^−11^ m^2^/s is adopted for this nanocarrier property in viable skin tissues. Relative to the base values in **Table 2**, the change in diffusion coefficients of nanocarriers in the skin cavity is by the same factor as the change in diffusion coefficient in the skin tissues in the following parametric simulations.

**Figure 8** compares the delivery results when using nanocarriers with different diffusion coefficients in the skin. Reducing this diffusion coefficient can slow down the nanocarrier escape from the patch since the decelerated nanocarrier transport into the tissues raises the concentration gradient across the patch-cavity interface. The peak of nanocarrier concentration varies greatly among the cavity, viable epidermis, papillary dermis and blood, presenting a non-linear response to the change in this property. However, the concentration in the reticular dermis is positively related to this diffusion coefficient. This is because rapid diffusion enables the nanocarriers to quickly transfer from the cavity to the surrounding skin tissues and then spread down to deeper skin layers. The concentration of free drugs and nanocarriers share similar trends in each layer and blood as shown in the middle panel. Comparisons of exposure to drugs over time demonstrate that the most effective treatment in the viable epidermis can be achieved when the nanocarrier diffusion coefficient in the skin tissue is around 1.0 × 10^−13^ m^2^/s . The optimum is 1.0 × 10^−14^ m^2^/s for the blood and papillary dermis. To be different, the effectiveness in the reticular dermis improves with this nanocarrier property.

**Figure 8.**
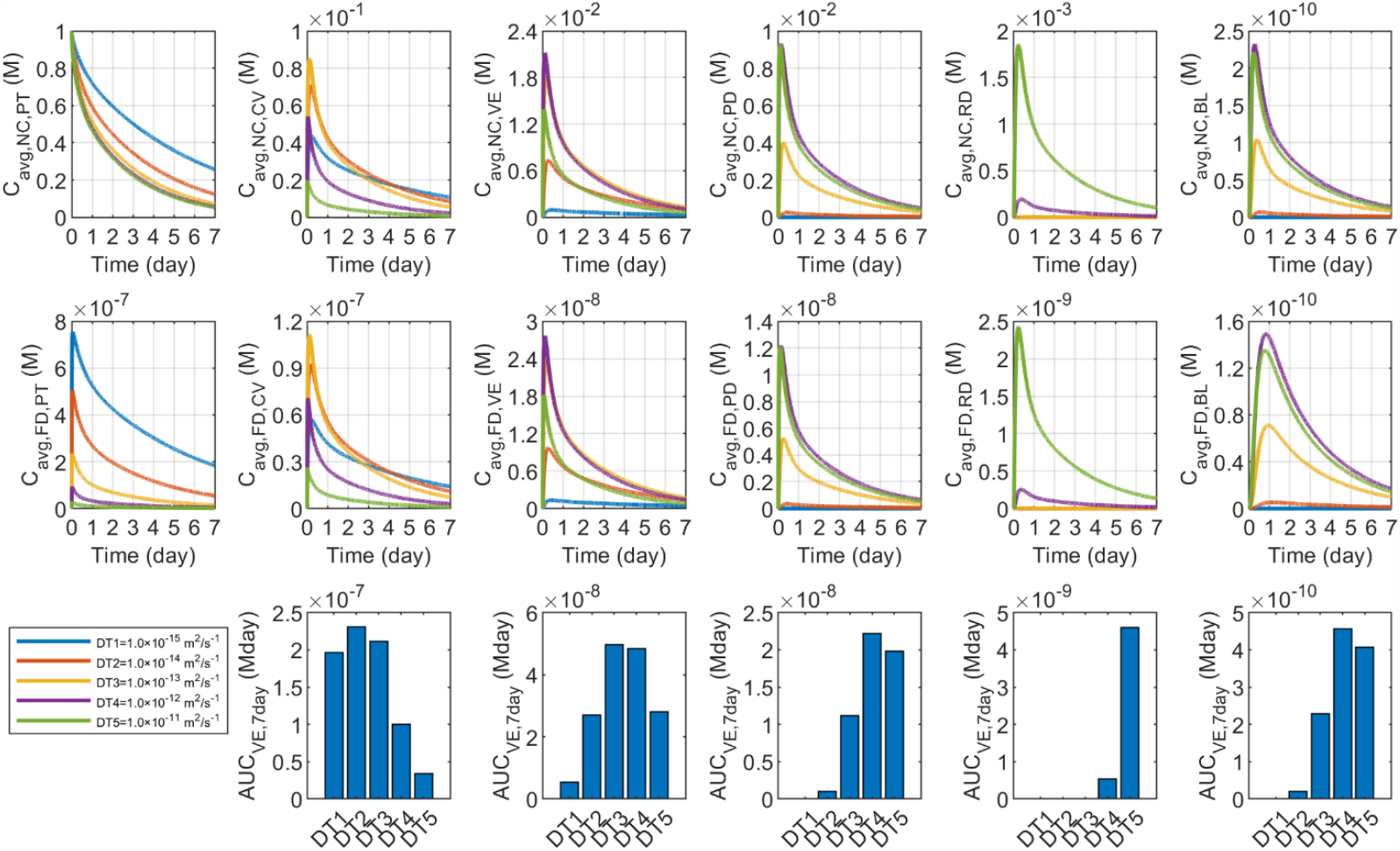
Impact of nanocarrier diffusion coefficient in the skin (*D*_NC,tis_) on the transdermal delivery results using solid microneedles.

#### 3.3.3 Vascular permeability

Vascular permeability refers to the ability of nanocarriers to pass through blood capillary walls. Large molecules are usually difficult to enter the bloodstream from the capillaries in the skin because of the tight connections between the endothelium cells on the capillary walls. However, this transport can be enhanced by modifying the nanocarrier formulation and size. Previous experiments have reported that permeability changes are on the order of 10^−11^ - 10^−9^ m/s [25]. Theoretical analysis further showed that reducing the drug carrier’s dimension to a few nanometres could raise this parameter to the order of 10^−7^ m/s [26]. Therefore, the range of 1.0 × 10^−11^-1.0 × 10^−7^ m/s is used. The impact of nanocarrier vascular permeability on drug delivery results is shown in **Figure 9**. Drug concentration in the patch, cavity and viable epidermis slightly decreases with this parameter. In contrast, the impact on the nanocarrier concentration is more pronounced in the papillary dermis, since blood capillaries are mainly embedded in this layer. This enhanced transport across vessel walls thereby increases free drug blood concentrations, however, the free drug deposition in the papillary and reticular dermis is reduced since fewer nanocarriers are left in these two layers to release the payload. Moreover, the exposure to drugs, *AUC*, decreases with increased vascular permeability in all the skin layers. Correspondingly, *AUC* in the blood rises sharply with this permeability, particularly when it is greater than 1.0 × 10^−9^ m/s.

**Figure 9.**
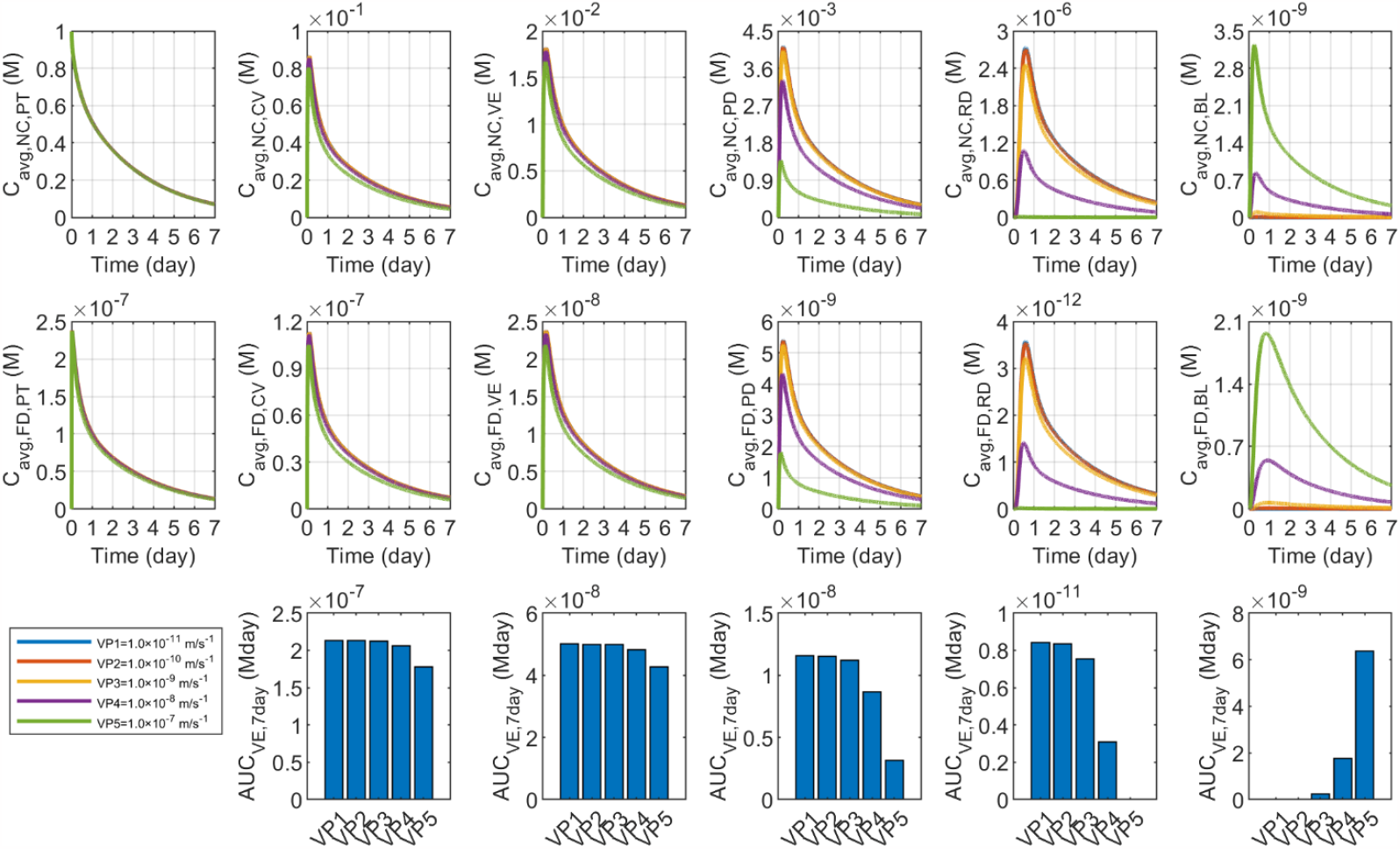
Impact of nanocarrier vascular permeability (*P*_NC_) on the transdermal delivery results using solid microneedles.

### 3.4 Role of patch properties

#### 3.4.1 Patch thickness

The thickness of the medicated adhesive patch can be well controlled in fabrication. This parameter was measured as 100 μm in Ref.[8], while a 250 μm thick patch was used with solid microneedles in Ref.[27]. The range of 40 - 280 μm is applied to determine its impact. Please note that the initial nanocarrier concentration in the patch changes simultaneously with the patch thickness to enable the identical dose for administration.

The delivery results of patches with different thicknesses are displayed in **Figure 10**. The nanocarrier concentration is found to decrease more rapidly in a thin patch. This is because the higher initial concentration generates a sharper concentration gradient across the patch-cavity interface; moreover, the shorter transport distance in the patch enables more efficient diffusion from the patch to the cavity. Consequently, the nanocarrier concentration rises faster to a higher peak in all the tissue compartments. However, due to the less durable drug supply from the thin patch, this concentration also decreases rapidly to a relatively lower level as time proceeds. The free drug concentration in the blood and each skin layer exhibit a similar response to patch thickness corresponding to the local nanocarrier concentration. The lower panel shows that the therapeutic effect decreases with increasing patch thickness when the dose is held constant.

**Figure 10.**
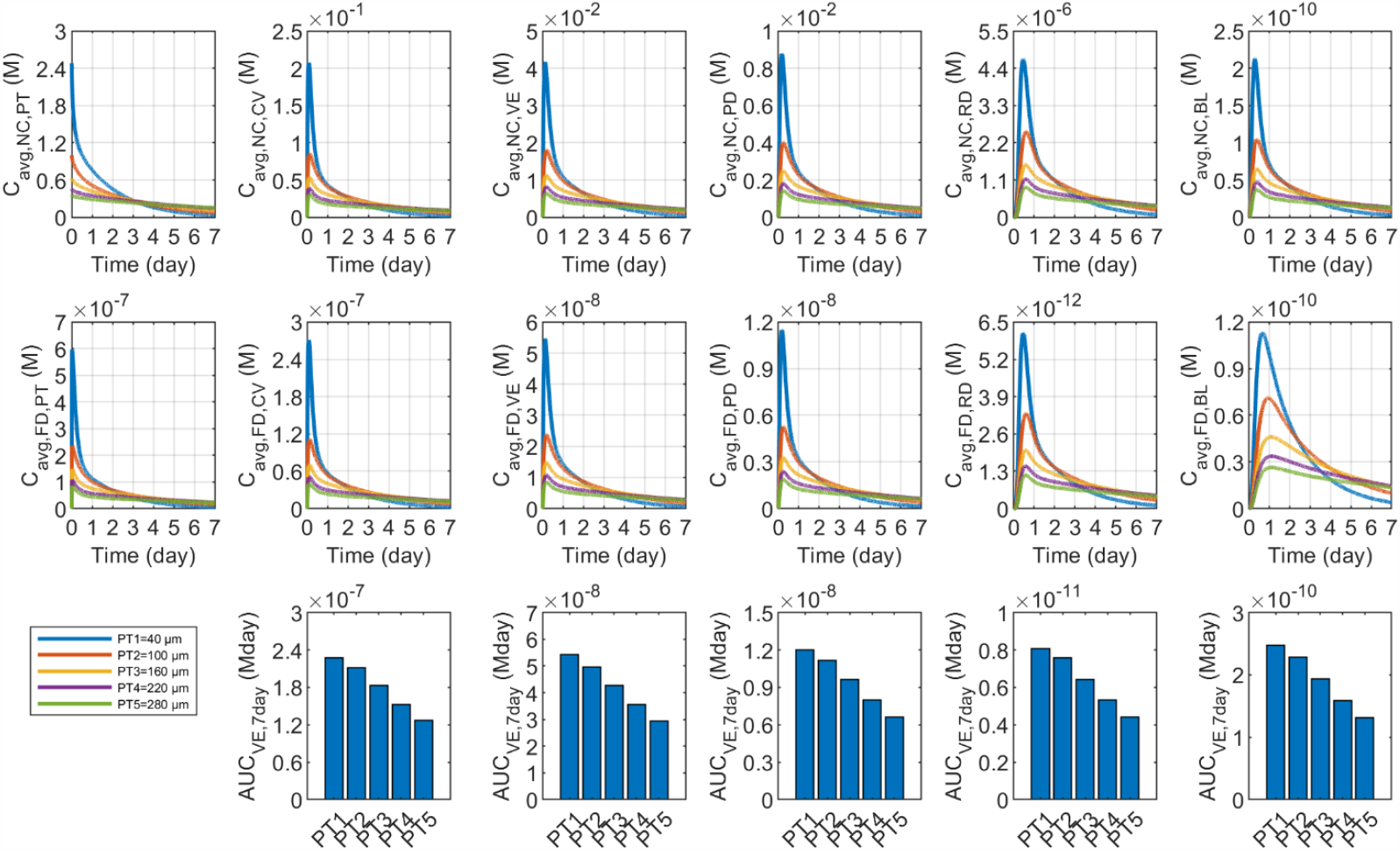
Impact of patch thickness (*h*) on the transdermal delivery results using solid microneedles.

#### 3.4.2 Diffusion coefficient of nanocarriers in patch

This diffusion coefficient is highly related to the physicochemical properties of the patch, especially the materials. Since this parameter was measured at the scale of 10^−13^ to 10^−12^ m^2^/s in polymers [28] and 10^−13^ to 10^−11^ m^2^/s in hydrogels [29], respectively, the range from 1.0 × 10^−15^ to 1.0 × 10^−11^ m^2^/s is used. The influence of this diffusion coefficient on the delivery results is shown in **Figure 11**. The nanocarriers which diffuse faster in the patch can travel more efficiently to the patch-cavity interface, thereby accelerating the drug delivery to the skin. As a result, the nanocarrier concentration can reach a higher peak in all the tissue compartments and blood but decreases faster to lower levels because of the less stainable supply of drugs from the patch. A similar response of free drugs can be found in the cavity, skin and blood, reflecting the direct influence of nanocarriers on free drugs. The lower panel of **Figure 11** shows a positive relationship between the treatment and this nanocarrier property in the blood and all skin layers, however, the increase in *AUC* becomes less pronounced when this diffusion coefficient is higher than 1.0 × 10^−12^ m^2^/s.

**Figure 11.**
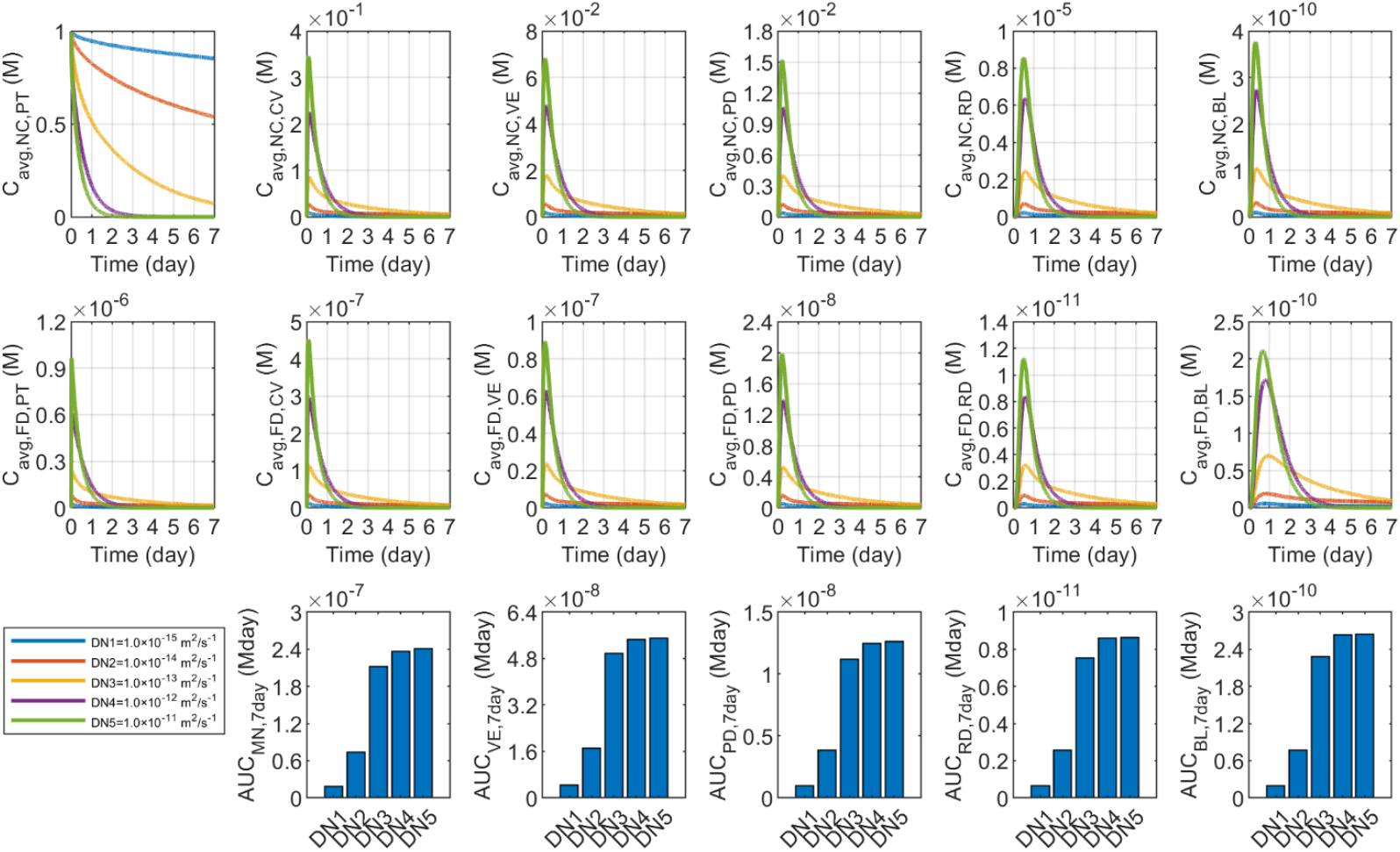
Impact of nanocarrier diffusion coefficient in the patch (*D*_NC,PT_) on the transdermal delivery results using solid microneedles.

#### 3.4.3 Nanocarrier partition coefficient

The partition coefficient represents the distribution ability of nanocarriers in different media. This parameter was found to lie in the range of 0.07 - 27.78 when the drug nanocarriers were dispersed in water and octanol [30,31]. The base value is 1.0 based on the assumption that the patch is aqueous. A large range of 1.0 × 10^−2^ - 1.0 × 10^2^ is then applied to examine its role. The case where this coefficient equals 0 represents the fully dispersed nanocarriers in the patch.

Shown in **Figure 12** are the responses of delivery results to the nanocarrier partition coefficient. The drug concentrations in all the tissue compartments are maintained at zero over time in the extreme case since no drug can escape the patch. Due to the enhanced distribution ability of nanocarriers in tissue fluid, the concentration of nanocarriers in the patch decreases rapidly as the partition coefficient increases, leading to a sharp rise in the concentration in all the tissue compartments. Notably, although the nanocarrier concentration reaches a higher peak as the partition coefficient increases, it decreases quickly to a lower level as time proceeds. The concentrations of free drugs and nanocarriers share similar trends in each compartment except the patch. This is because nanocarriers well encapsulate free drugs before travelling to the skin. So that all free drugs in the patch are from the cavity. The lower panel shows the treatment in the blood and all the skin layers can be improved by raising the nanocarrier partition coefficient. However, this improvement becomes less significant when this parameter is greater than 1.0.

**Figure 12.**
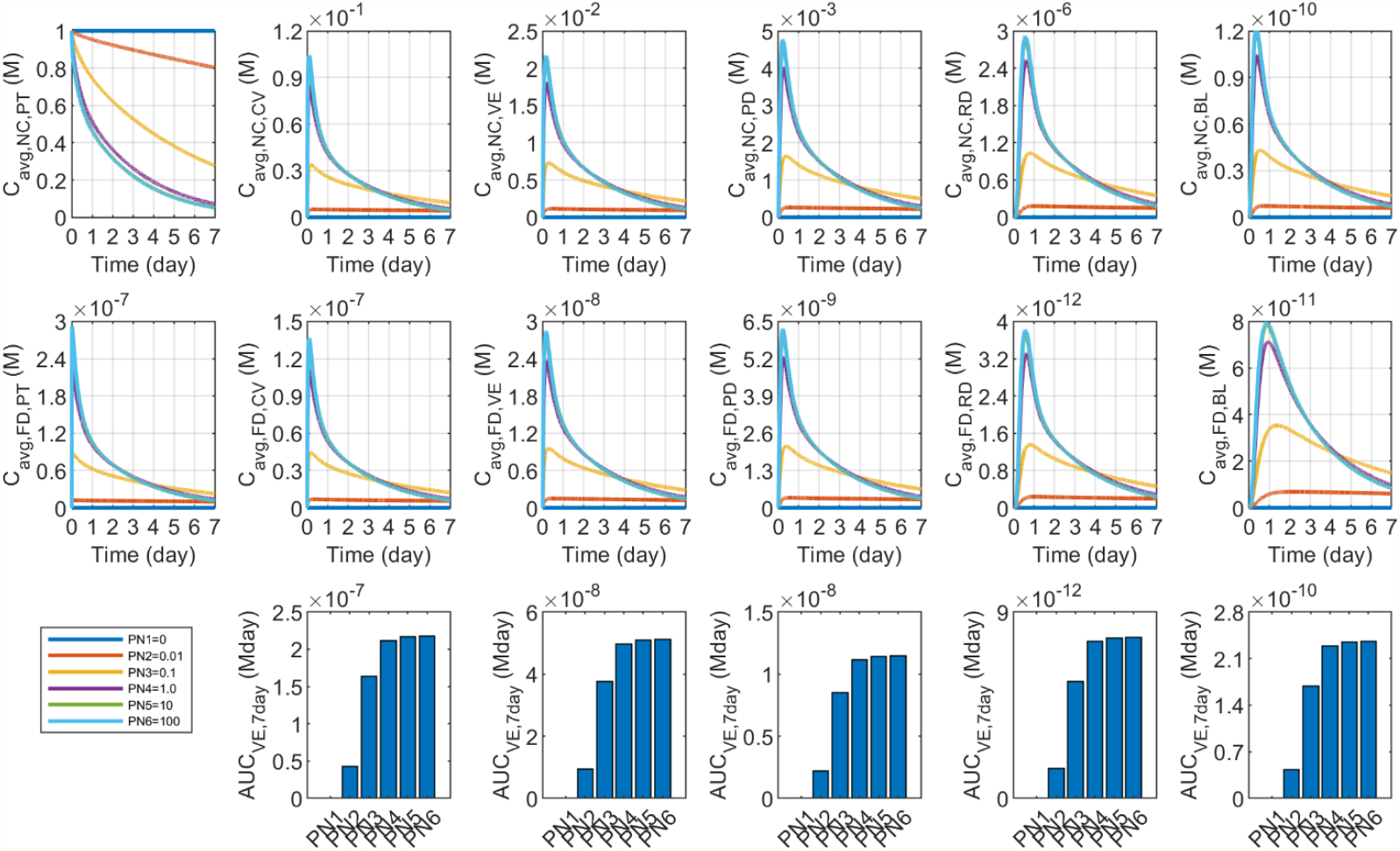
Impact of nanocarrier partition coefficient (*K*_NC_) on the transdermal delivery results using solid microneedles.

The mathematical model is also applied to determine the impacts of patch-induced changes in free drug properties, including the physical degradation rate and diffusion coefficient of free drugs in the patch, and the free drug partition coefficient between the cavity and patch. The results given in the supplementary document demonstrate that the delivery results in the skin and the blood circulatory system are less sensitive to these parameters.

### 3.5 Impact of microvasculature

#### 3.5.1 Blood capillary density

The density of blood capillaries reflects the blood vessel distribution in the papillary dermis. It would directly affect the amount of drugs transferring into the blood and thence systemic absorption. This tissue property varies subject to the location in the human body and between patients. It is described by *S*_bl_/*V*_tis_ in this mathematical model, but its value in the skin is rarely reported in the literature. A representative value is calculated as 10550 m^−1^ given the average surface area of capillary walls in the entire human body and the tissue density were reported to be 10 m^2^/kg [32] and 1055 kg/m^3^ [33], respectively. The surface area of blood capillaries in the skeletal muscle is usually considered two times higher than that in the skin [34]. Since the value in skeletal muscle was measured as 8.4 m^2^/kg [32], the *S*_bl_/*V*_tis_ in the skin is estimated to be 4431 m^−1^ . In order to cover the possible values of this tissue property, the range from 4000 to 12000 m^−1^ is adopted in this study.

**Figure 13** shows the results of transdermal delivery to the skin with different blood capillary densities. Administrating drugs to the capillary-rich skin can effectively accelerate the increase in the blood concentrations of both free drugs and nanocarriers to higher peaks since more drugs can cross the capillary walls; these concentrations in this skin can also sustain at relatively higher levels over time to provide a more durable drug supply to benefit systemic absorption. Moreover, this tissue property has a neglectable influence on drug deposition in the upstream compartments, including the patch, cavity and viable epidermis. Drug concentrations in the papillary dermis and reticular dermis slightly decrease due to the reduced drug availability in these two skin layers. Consequently, drug exposure in the blood presents a positive relationship with the blood capillary density, whereas the treatment in the papillary dermis and reticular dermis decreases.

**Figure 13.**
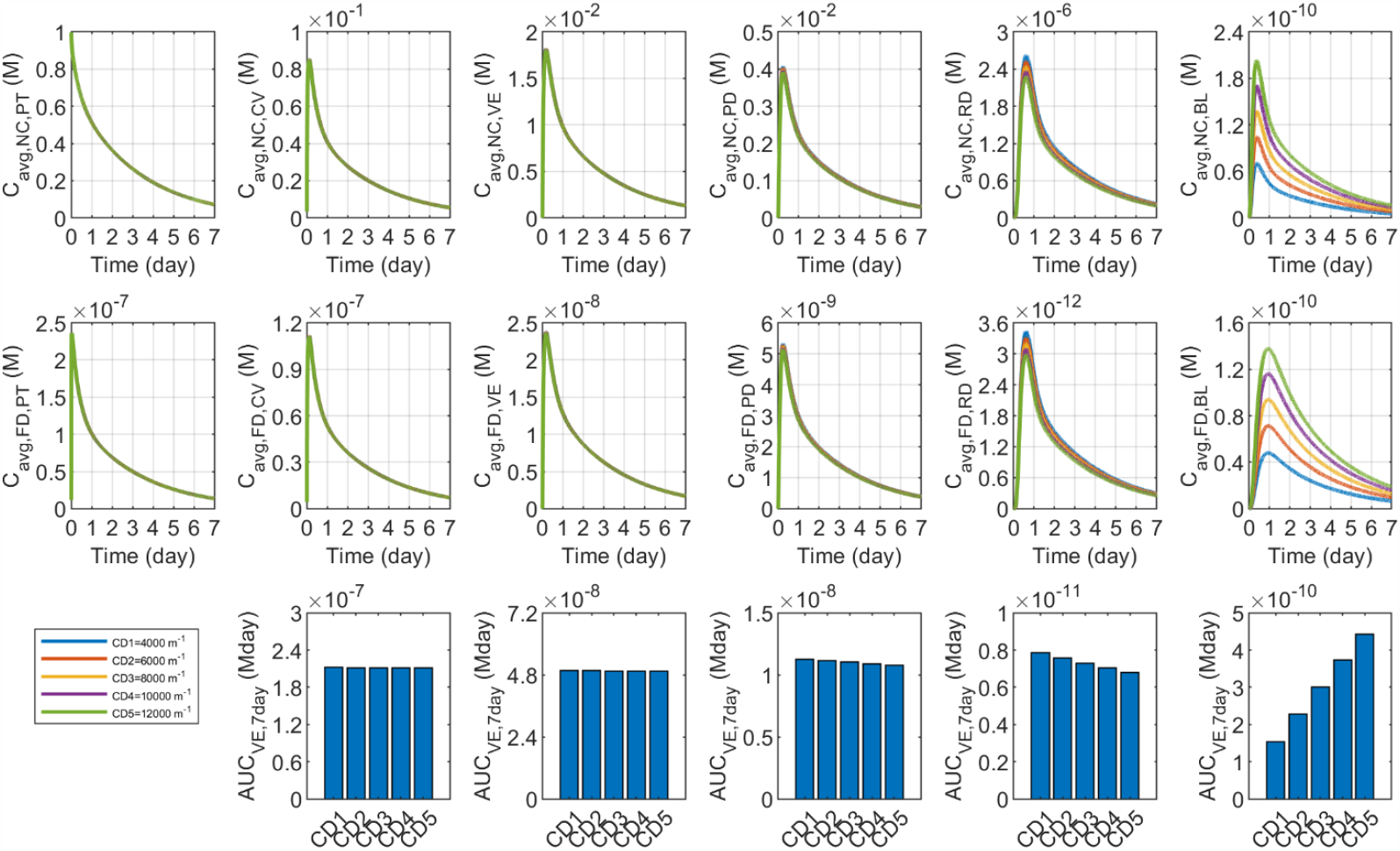
Impact of blood capillary density (*S*_bl_/*V*_tis_) on the transdermal delivery results using solid microneedles.

#### 3.5.2 Lymphatic function

The lymphatics can effectively remove the drugs from the tissue due to the highly permeable nature of the lymphatic walls. Since this function has been found to decline with age [35], the full function in **Table 2** is reduced to 75%, 50%, 25% and 0 in this study to reveal its role. Results in **Figure 14** demonstrate that the time courses of drug concentration and drug exposure are not sensitive to the change in this tissue property, indicating a neglectable impact of lymphatic drainage on the outcomes of this transdermal drug delivery.

**Figure 14.**
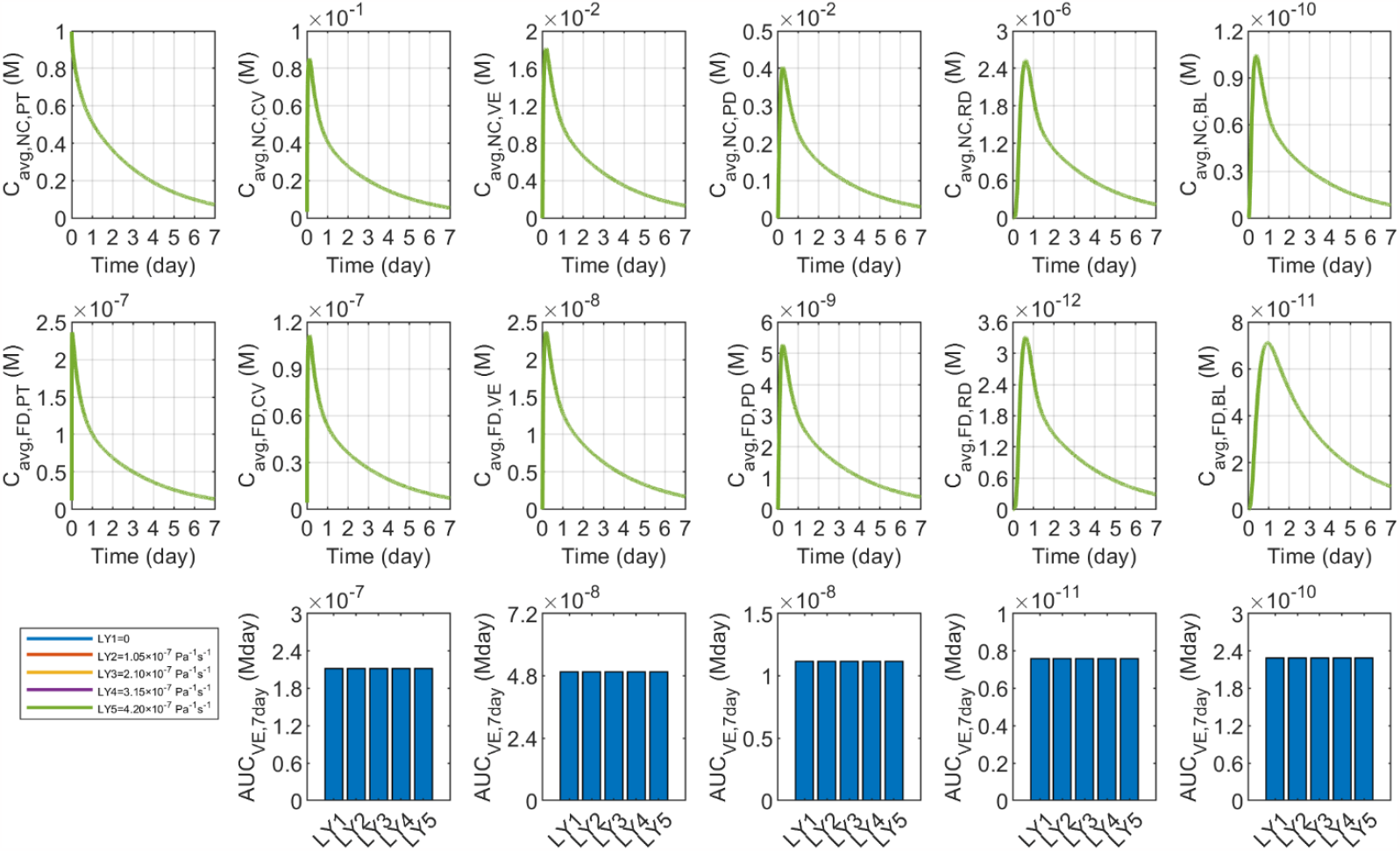
Impact of lymphatic function (*L*_ly_*S*_ly_/*V*_tis_) on the transdermal delivery results using solid microneedles.

## 4. Discussion

Stratum corneum is the major obstacle to transdermal drug delivery owing to its nearly impermeable nature. Microneedles can effectively pierce the stratum corneum and thereby enable drugs to be delivered to viable skin tissues and circulatory systems. The combination of solid microneedles and medicated adhesive patches can provide longer-lasting treatment compared to the microneedles which load drugs inside [12]. Moreover, drug delivery results differ greatly across skin layers and blood in response to changes in drug delivery conditions, as demonstrated by the simulation results, stressing the need to tailor the drug delivery system to meet the particular treatment requirement. For instance, delivery to a specific layer can be improved by careful selection of nanocarriers with optimal release rate or microneedle length enabling the microneedle tip to reach that layer. Increasing the nanocarrier vascular permeability or delivering drugs to capillary-rich skin area is effective in improving drug delivery to the blood circulatory system, and simultaneously reducing the therapeutic effect on all skin tissues. To be different, reducing the patch thickness, microneedle spacing or raising the nanocarrier diffusivity in the patch can successfully enhance the treatment in the blood and all skin layers.

The applied mathematical model in this study predicts drug transport and deposition in different skin layers and circulating systems by simulating the interplays between the tissue and therapeutic agents. The simulation predicted concentrations of desoximetasoneas as a function of the depth in the skin tissue is compared with experimental results in **Figure 15** under the same delivery condition as in Ref.[36]. Similar model validation has been reported in studies using mathematical models to predict transdermal and topical drug delivery [12,36] and drug delivery to different tissues [37]. However, it is still important to point out that numerical simulations can usually provide qualitative predictions on drug delivery outcomes. This is mainly because a comprehensive model capable of describing all drug-tissue interactions at sufficiently high resolution is still lacking, and the simulation needs a large number of model parameters which can vary greatly depending on time, location of the lesion and patients. However, acquiring all these patient-specific parameters remains less feasible. Despite these limitations, the qualitative predictions are sufficient for revealing the impacts of influencing factors through cross-comparisons, allowing for identifying opportunities to improve drug delivery systems and delivery strategies for better treatment.

**Figure 15.**
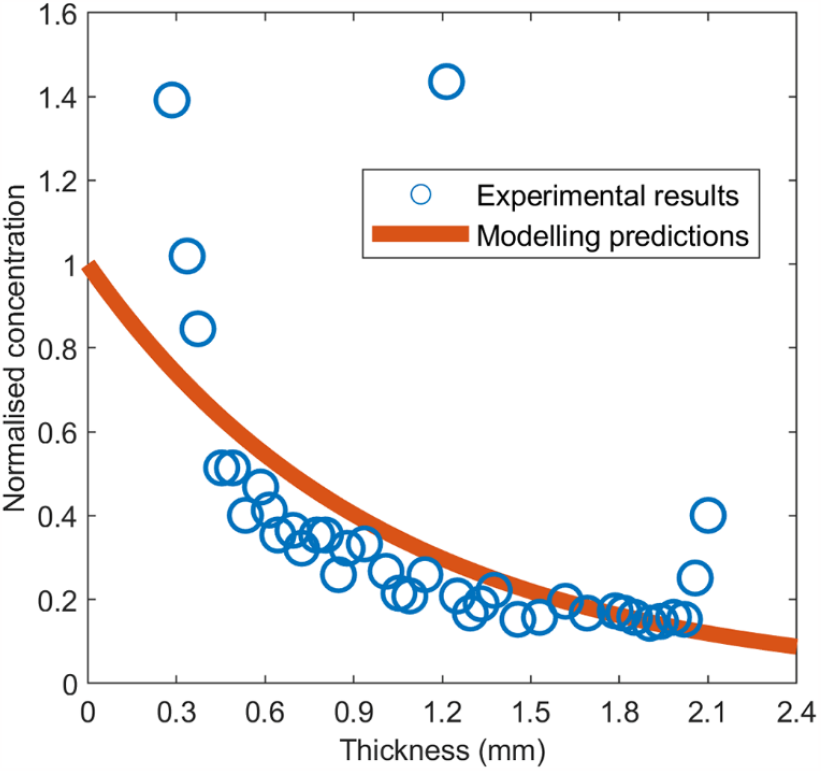
Comparison of the concentration of desoximetasone as a function of penetration depth in the skin between simulation predictions and experimental measurements. The experimental results and drug properties are obtained from Ref. [36]. The coefficient of determination is calculated as 0.65.

Nevertheless, several limitations are involved in this study. (1) The cavity is assumed to have the same morphologic features as the solid microneedle. In practice, the cavity boundaries can become irregular after the microneedles are withdrawn, due to the soft nature of skin tissues. Microscopic images of the treated skin tissues can be used to reconstruct the real geometry of the cavity to relax this assumption. (2) The water molecule-impermeable patch is studied in the simulations. However, the water loss through the drug-loaded patch may take place, depending on the patch materials and environment. This process is not included in this study due to the lack of a validated model to describe water transport in the patch and escape into the atmosphere. This limitation highlights the need for developing the model to accommodate more comprehensive delivery conditions. (3) The delivery results also highly depend on the delivery regime, such as the administration dose, duration of the patch placed on the skin surface, treatment cycles and dosing intervals. The influences of these factors are not examined in this study. For instance, the patch stays on the skin surface for a week to enable an enough large time window to monitor the continuous change in drug concentration in different tissue compartments. A future study can be performed to optimise delivery regimes and further develop the strategy for this transdermal delivery.

## 5. Conclusions

Transdermal delivery of drug nanocarriers using solid microneedles has been studied under different delivery conditions. The numerical simulations demonstrate the dominance of diffusion in determining the nanocarrier and free drug transport in the skin. Delivery outcomes differ distinctly in response to the changes in the properties of the medicated adhesive patch, solid microneedle, nanocarriers and microvasculature. Specifically, the drug release rate needs to be selected concerning the depth of the target site in the skin to maximise the treatment. Drug exposure in the blood and entire skin exhibits a positive relationship with the partition coefficient and diffusion coefficient of nanocarriers in the patch; however, optimisation is needed for the property of nanocarrier diffusion coefficient in the skin to improve treatment for the viable epidermis, papillary dermis and blood. Improving the vascular permeability of nanocarriers or placing solid microneedles in areas of skin with dense capillaries can effectively enhance drug exposure in the blood circulatory system while reducing therapeutic efficacy in all skin tissues. The treatment for a specific layer can be effectively improved by using microneedles with an appropriate length to position the microneedle tip at that layer. Moreover, using a thin medicated adhesive patch or closely packed microneedle array can improve treatment outcomes in the skin and blood for a given injected dose. However, the delivery outcomes are less sensitive to the transport properties of free drugs in the patch. These findings can serve as a reference to improve transdermal drug delivery outcomes using solid microneedles and medicated adhesive patches.

## Notes

### Competing Interest Statement

The authors have declared no competing interest.

