## Supplemental figures for "Numerical Simulation of Transdermal Delivery of Drug Nanocarriers Using Solid Microneedles"

The mathematical model is applied to examine the free drug properties in the patch. The results are summarised below.

1. Partition coefficient of free drugs

This partition coefficient evaluates the distribution ability of the representative drug in the patch and interstitial fluid in the cavity, subject to the properties of drug molecules, interstitial fluid and patch. The reported partition coefficient of doxorubicin between water and octanol was $20.8$ [1] and $17.9$ [2]. $1.0$ is selected as the base value by assuming the patch is an aqueous phase. The range from $0$ to $100$ is used to cover the possible levels this property can arrive at. The case of $0$ stands for the patch which is perfectly dispersible for the free drugs.

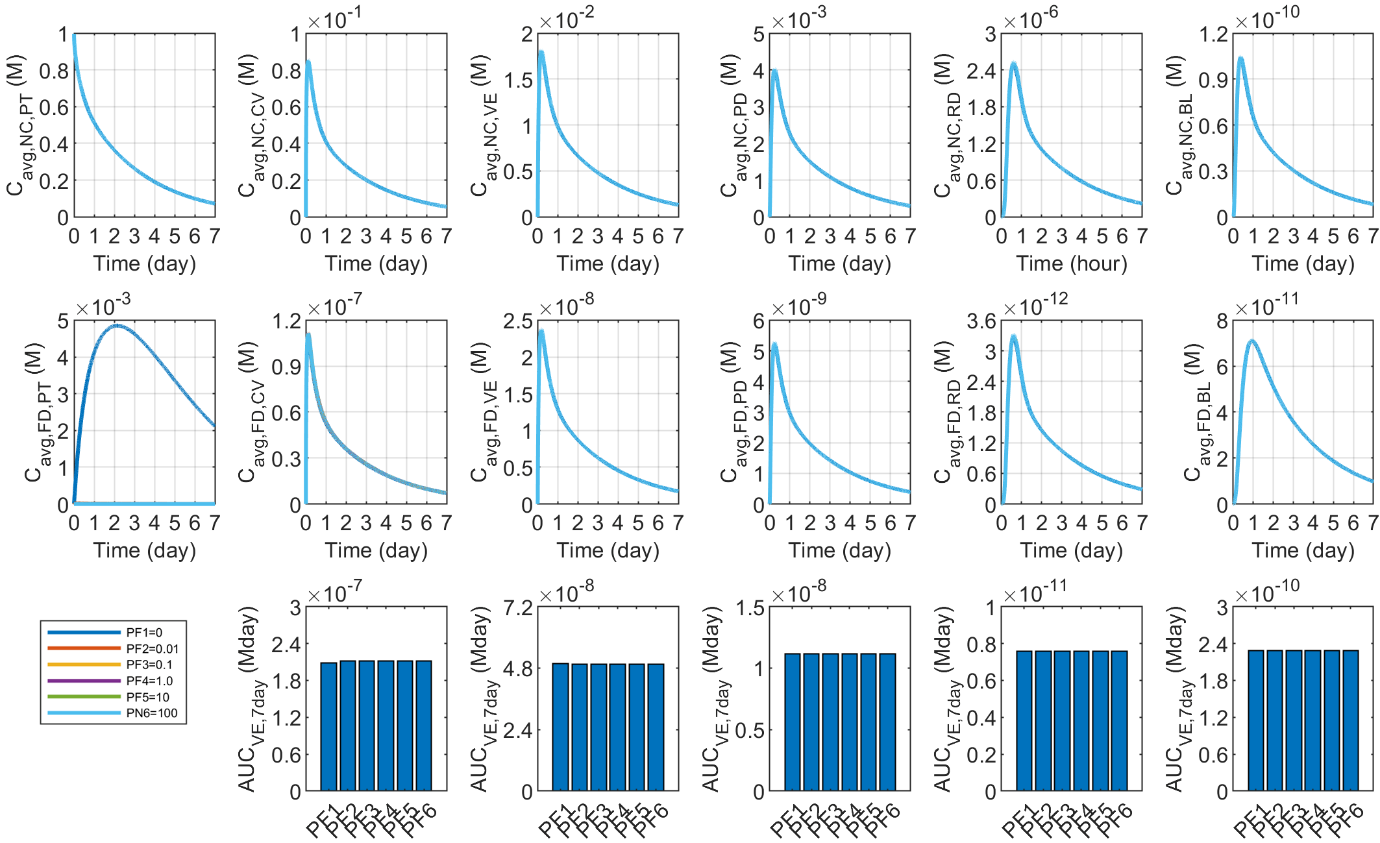

Figure S1. Impact of free drug partition coefficient in the patch ($K_{\mathrm{FD}}$) on the transdermal delivery results using solid microneedles.

**Figure S1** summarises the response of drug delivery results to this partition coefficient. The simulations show that there is no obvious difference in the concentration of nanocarriers between each skin layer and the blood when this property changes. Although reducing this partition coefficient to an extremely low level can greatly raise the free drug concentration in the medicated adhesive patch, the influence on the time course of free drug concentration and drug exposure in the tissue compartments is neglectable.

2. Diffusion coefficient of free drugs in patch

This diffusion coefficient describes the ability of drug molecules to travel in the patch due to thermal motion. Its value in the patch is strongly determined by the patch materials. Since the diffusivity of doxorubicin in hydrogels and polymers was measured as $8.2\times{10}^{-10}$ - $2.3\times{10}^{-9} {m^{2}}/s$ [3] and $1.9\times{10}^{-11}$ - $1.2\times{10}^{-10} {m^{2}}/s$ [4], respectively, the range from $1.0\times{10}^{-12}$ to 1.0$\times{10}^{-8} {m^{2}}/s$ is adopted to determine its influence. The results in **Figure S2** demonstrate that raising this parameter can slightly increase the peak concentration of free drugs in the patch. However, delivery results in the skin and blood remain almost unchanged.

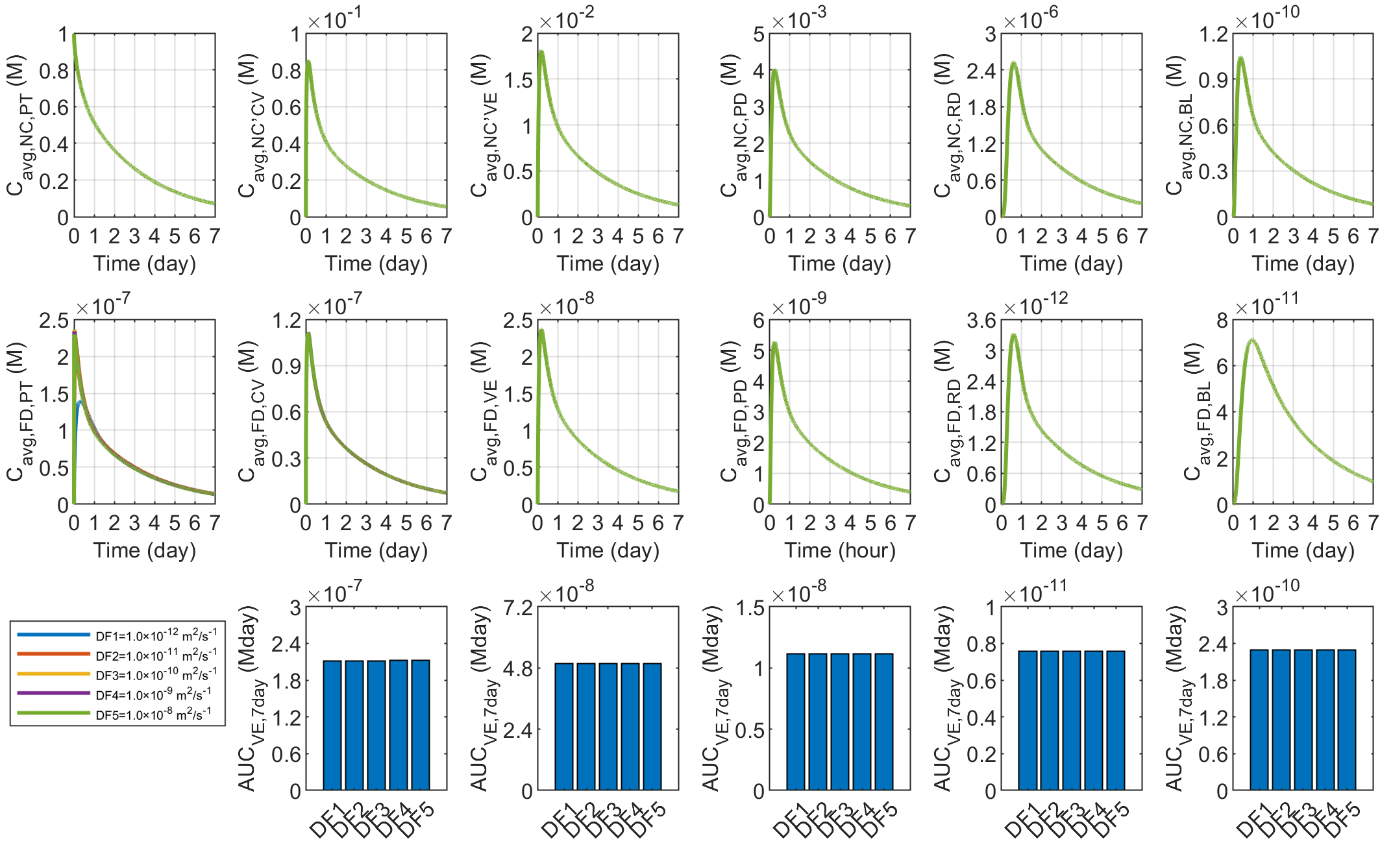

Figure S2. Impact of free drug diffusion coefficient in the patch ($D_{FD,PT}$) on the transdermal delivery results using solid microneedles.

3. Physical degradation rate of free drugs in patch

The rate of physical degradation reflects how fast drugs are eliminated owing to the change to materials in the surrounding environment, influenced by the patch materials. For instance, doxorubicin degrades slower in polymers than in phosphate-buffered saline [5]. Therefore, the base value of the degradation rate in **Table 2** is changed from $0.01$ to $100$ times to conduct an exploratory parametric study. As presented in **Figure 14**, the drug concentration and exposure remain almost constant as this rate changes, indicating the very limited influence of the drug's physical degradation rate on the delivery outcomes.

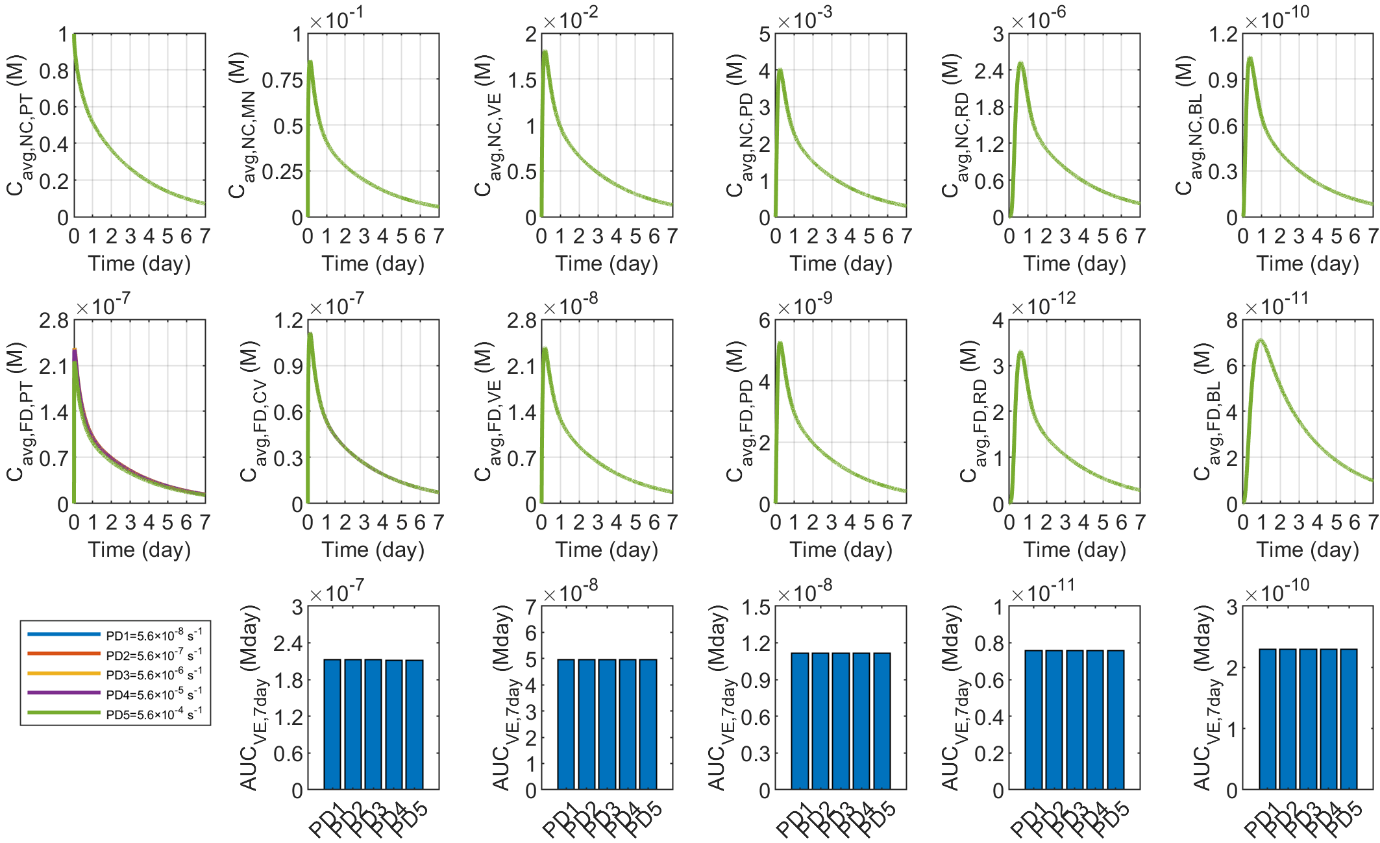

Figure S3. Impact of free drug degradation rate in the patch ($k_{d,PT}$) on the transdermal delivery results using solid microneedles.
